# Molecular and functional profiling distinguishes PACS1 syndrome variant from PACS1 loss-of-function in iNeurons

**DOI:** 10.64898/2026.08.25.747101

**Authors:** Annika L Schroder, Ximena Gomez-Maqueo, Sean R Golinski, Claire M Phoumyvong, Richard S Smith, Alicia Guemez-Gamboa

## Abstract

PACS1 syndrome is a rare neurodevelopmental disorder caused by a recurrent *de novo* missense variant (p.R203W) in the PACS1 protein. However, it remains unclear whether the p.R203W variant acts through a loss-of-function or alternative mechanism. Here, we used isogenic iPSC-derived neurons (iNs) to directly compare the effects of PACS1 p.R203W to complete loss of PACS1 function. Using a combination of proteomic, biochemical and electrophysiological approaches, we identified molecular and functional phenotypes associated with each genotype. While PACS1^(+/R203W)^ and PACS1^(-/-)^ iNs shared phenotypic abnormalities, the overall molecular and functional consequences of the p.R203W variant were distinct from those caused by PACS1 deficiency. Notably, PACS1^(+/R203W)^ presented with unique proteomic and kinase signaling signatures and a shift in stimulus dependent excitability. These findings demonstrate that PACS1 syndrome is not caused by a simple loss of function and instead support a non-loss-of-function mechanism. Lastly, our interactome analysis suggests that the p.R203W variant retains aspects of canonical PACS1 function while acquiring novel molecular interactions that could contribute to PACS1 syndrome pathogenesis. Altogether, these findings provide a framework for future mechanistic studies and therapeutic development in PACS1 syndrome.

**GRAPHICAL ABSTRACT:** 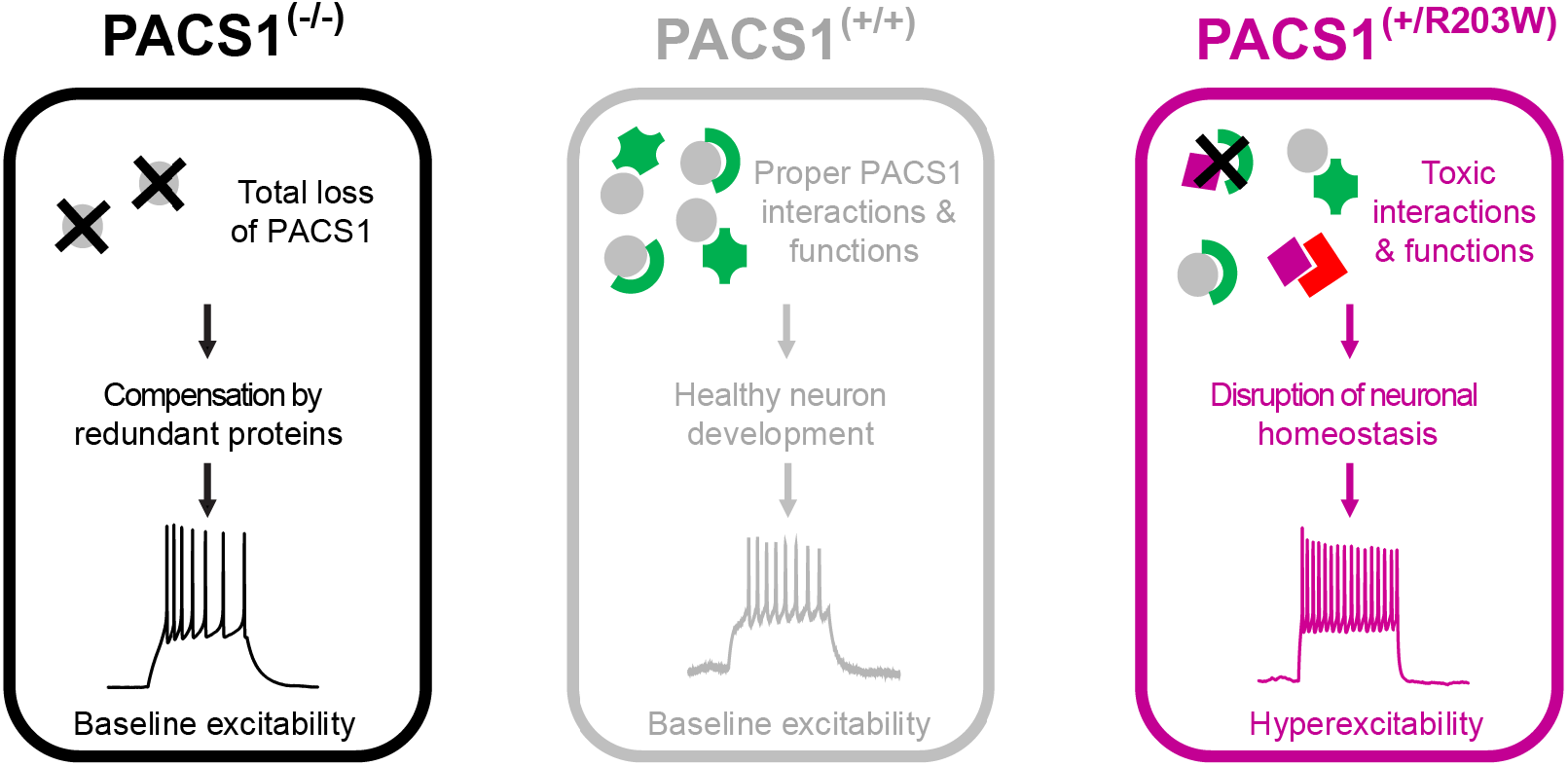

## INTRODUCTION

Neurodevelopmental disorders (NDDs) are a group of disorders with high degrees of genetic and clinical variability that affect the early development of the cerebral cortex, often impairing the ability to reach cognitive and behavioral milestones (1, 2). Monogenic NDDs are caused by variation of single genes and can be caused by *de novo* missense variants, meaning they are not inherited from either parent (3). It is critical to understand how these *de novo* pathogenic variants influence neurodevelopment, whether that be through a loss-of-function (LoF), gain-of-function (GoF) or dominant negative mechanism, as this will inform therapeutic intervention to safely alleviate symptoms of affected individuals.

PACS1 syndrome is an example of a monogenic NDD that is caused by a recurrent *de novo* missense variant (c.607C>T) in the Phosphofurin acidic cluster sorting 1 (*PACS1*) gene, resulting in a p.R203W substitution in the PACS1 protein (4–13). Individuals with PACS1 syndrome have an unaffected allele in addition to the p.R203W variant. The PACS1 syndrome phenotype is variable, but all individuals present with intellectual disability and craniofacial dysmorphisms, and nearly half have a diagnosis of epilepsy (4–14). Additionally, as NDDs often converge in molecular pathways related to chromatin remodeling and regulation of gene expression, it is important to note that a mild epigenetic signature is associated with PACS1 syndrome (15–18).

PACS1 is a multifunctional sorting protein canonically described for its role in regulating trans-Golgi Network (tGN) and endosome vesicle trafficking. PACS1 was first discovered for its interaction with Furin and is required for its localization to the tGN (19, 20). Since then, it has been shown that PACS1 interacts with a variety of proteins via acidic-cluster motifs (19, 20). PACS1 binding partners now extend beyond regulators of tGN and endosomal trafficking (AP-1, AP-3, CI-MPR, Sortilin), and include chromatin remodelers (HDAC2, HDAC3), regulator of calcium in the endoplasmic reticulum WDR37, primary cilia components (NPHP1, TRPP2, CNGB1b, HDAC6), and synaptic proteins (SNAP25, VAMP4) (19–34). The identification of these binding partners has painted a picture where PACS1 is involved in much more than the tGN-endosomal pathways and has broadened how we think about PACS1 function in the cytoplasm and the nucleus.

Research investigating the impact of PACS1 p.R203W on neural development suggests it may have an impact on synapse formation and function (22, 35, 36). Work done in patient-derived iPSC forebrain organoids showed that mature glutamatergic neurons have impaired expression of genes associated with ion transport and synaptic signaling. Additionally, patient-derived organoids have an increased density of GABAergic synaptic markers (35). In mice, the overexpression of PACS1 p.R203W increases neurite arborization while reducing spine density. These neurons also display decreased amplitude and frequency of mEPSCs highlighting a potential decrease in overall synaptic strength at the pre and post synapse (22).

An essential part of understanding the pathogenic mechanism of the p.R203W variant is to understand whether the pathogenic variant behaves as a loss of PACS1 function. Previous studies investigating PACS1 p.R203W have not included PACS1 knockdown or knockout models, which has left this question unanswered (22, 35, 37). Current treatments for PACS1 syndrome are limited to N-of-1 trials of non-allele-specific antisense oligonucleotides (ASOs). These ASOs will lead to an overall decreased abundance of PACS1 in patients, which serves as another compelling reason to finally establish if there is a link between the p.R203W variant and loss of PACS1 function.

In this study we used isogenic induced pluripotent stem cells (iPSC)-derived neurons (iNs) to directly compare the effects of the recurrent p.R203W variant with loss of PACS1 function in an effort to address the limited understanding of PACS1 syndrome pathogenesis. Using proteomic, biochemical, and electrophysiological analyses, we demonstrate that while there are some convergent phenotypes between PACS1^(+/R203W)^ and PACS1^(-/-)^ iNs, PACS1 p.R203W ultimately produces molecular and functional phenotypes that are distinct from PACS1 loss of function, supporting a non-loss-of-function mechanism. Lastly, we identified variant-specific candidate interactors, suggesting that PACS1 p.R203W retains canonical PACS1 functions while obtaining pathogenic activities through the acquisition of novel molecular interactions that could contribute to disease pathogenesis. Collectively, our findings suggest that PACS1 syndrome is unlikely to result from simple loss of PACS1 function and instead support a non-loss-of-function mechanism for the pathogenic variant. By defining the molecular and cellular consequences of PACS1 p.R203W, this study provides a framework for future investigations into PACS1 syndrome pathogenesis and identifies candidate pathways that may inform therapeutic development.

## RESULTS

### Golgi and endosomal organization are preserved in PACS1^(+/R203W)^ and PACS1^(-/-)^ iNs

To understand if there is a relationship between PACS1 p.R203W and loss of PACS1 function, we differentiated isogenic PACS1^(+/+)^, PACS1^(+/R203W)^ and PACS1^(-/-)^ iPSCs into neurons (iNs) using a doxycycline-inducible Neurogenin2 (Ngn2) overexpression system (Figure 1A). We confirmed that PACS1 protein abundance did not differ between our PACS1^(+/+)^ and PACS1^(+/R203W)^ lines, which has been previously reported (Figure 1B-C) (35).

**Figure 1.**
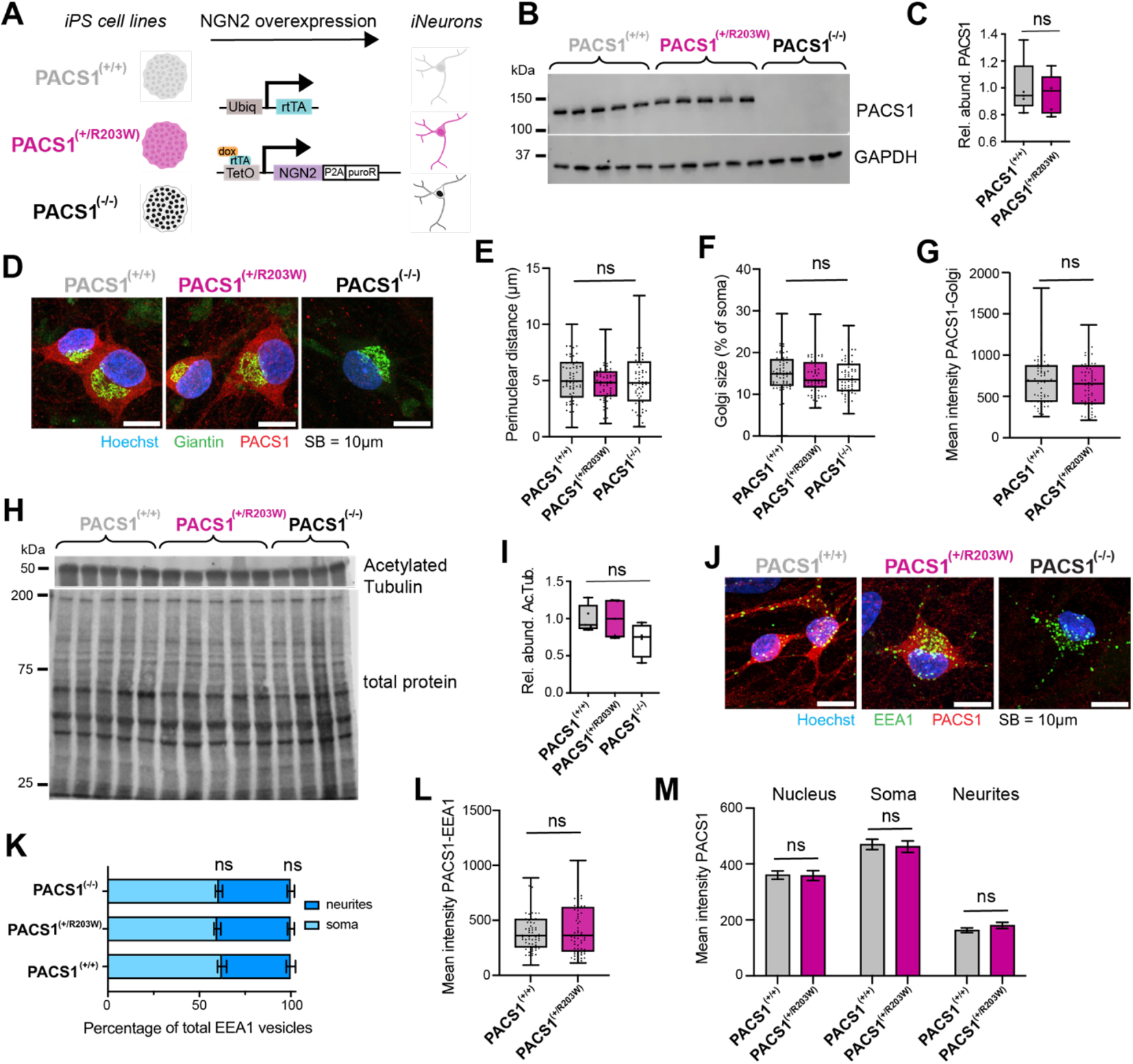
Golgi and endosomal organization are preserved in PACS1^(+/R203W)^ and PACS1^(-/-)^ iNs. **(A)** Experimental workflow to differentiate PACS1^(+/+)^, PACS1^(+/R203W)^ and PACS1^(-/-)^ iPSCs into iNs. **(B)** Western blot for PACS1 and GAPDH protein abundance in d29 iNs. **(C)** Quantification of the ratio of PACS1:GAPDH in PACS1^(+/+)^ and PACS1^(+/R203W)^ iNs (Two-tailed Mann-Whitney, p>0.9999, n=5 samples for each genotype). **(D)** Representative images of Golgi apparatus and PACS1 in PACS1^(+/+)^, PACS1^(+/R203W)^ and PACS1^(-/-)^ d35 iNs, scale bar (SB) = 10μm. **(E)** Quantification of perinuclear distance (Kruskal-Wallis with Dunn’s multiple comparisons; p_Kruskal_=0.5786; p_R203W_=0.8868, p_KO_>0.9999; n=70 cells per genotype) and **(F)** Golgi size normalized to soma size across the three genotypes (Kruskal-Wallis with Dunn’s multiple comparisons; p_Kruskal_=0.2555; p_R203W_=0.6567, p_KO_=0.3491; n=70 cells per genotype). **(G)** Mean fluorescence intensity of PACS1 in the Golgi apparatus in PACS1^(+/+)^ and PACS1^(+/R203W)^ iNs (Two-tailed Mann-Whitney, p=0.4357, n=70 cells for each genotype). **(H)** Western blot for acetylated tubulin and total protein stain in d29 iNs. **(I)** Quantification of the ratio of acetylated tubulin: total protein in iNs across all three genotypes (Kruskal-Wallis with Dunn’s multiple comparisons; p_Kruskal_=0.1841; p_R203W_>0.9999, p_KO_=0.1555, n=5 samples for PACS1^(+/+)^ and PACS1^(+/R203W)^ and n=4 samples for PACS1^(-/-)^ iNs). **(J)** Representative images of EEA1 positive vesicles and PACS1 in PACS1^(+/+)^, PACS1^(+/R203W)^ and PACS1^(-/-)^ d35 iNs; SB = 10μm. **(K)** Quantification of EEA1 vesicles in soma vs neurites across the three genotypes (Kruskal-Wallis with Dunn’s multiple comparisons; p_Kruskal_=0.5978; p_R203W_>0.9999, p_KO_>0.9999; n=70 cells per genotype). **(L)** Mean fluorescence intensity of PACS1 in EEA1 positive vesicles in PACS1^(+/+)^ and PACS1^(+/R203W)^ iNs (Two-tailed Mann-Whitney, p=0.7255, n=70 cells for each genotype). **(M)** Mean fluorescence intensity of PACS1 in nucleus, soma and neurites of PACS1^(+/+)^ and PACS1^(+/R203W)^ d35 iNs (3 independent two-tailed Mann-Whitney, p_nucleus_=0.5184, p_soma_=0.6121, p_neurites_=0.8588; n=140 cells per genotype).

Previous studies suggested that p.R203W indirectly destabilizes microtubules and leads to Golgi fragmentation and dispersal (22, 31). To determine if PACS1^(+/R203W)^ iNs recapitulate this phenotype and establish a baseline comparison to PACS1^(-/-)^, we examined Golgi organization in day 35 iNs (Figure 1D). Contrary to previous reports, Golgi architecture was preserved across genotypes. Particularly, no differences were observed in Golgi perinuclear positioning, percentage of soma occupied by the Golgi, and PACS1 enrichment in the Golgi between PACS1^(+/+)^ and PACS1^(+/R203W)^ iNs (Figure 1F-G).

As the lack of a Golgi phenotype was unexpected, we examined microtubule stability by comparing abundance of acetylated tubulin between genotypes (Figure 1H). While previous studies reported reduced acetylated tubulin in PACS1^(+/R203W)^ cells, we observed comparable levels in PACS1^(+/+)^ and PACS1^(+/R203W)^ iNs. In contrast, PACS1^(-/-)^ iNs presented with a modest decrease in acetylated tubulin abundance (Figure 1I).

As PACS1 is canonically described as a regulator of endosomal trafficking, we next examined endosomal vesicle organization and PACS1 localization to early endosomes (Figure 1J). Again, we observed no differences in the subcellular distribution of EEA1 vesicles across genotypes, and that PACS1 enrichment within these vesicles was comparable between control and PACS1^(+/R203W)^ iNs (Figure 1K-L). Lastly, given the absence of Golgi or early endosome related phenotypes, we assessed the overall localization of PACS1. Quantification of PACS1 fluorescence revealed no differences in protein abundance in the nucleus, soma or neurites of PACS1^(+/+)^ and PACS1^(+/R203W)^ iNs (Figure 1M).

Altogether, these findings highlight that PACS1 p.R203W does not induce aberrant Golgi positioning or endosomal distribution phenotypes and does not affect PACS1 localization in our human neuronal model. Because we observed no major alterations in the canonical trafficking functions of PACS1, we next performed unbiased proteomics analysis of PACS1^(+/+)^, PACS1^(+/R203W)^ and PACS1^(-/-)^ iNs to identify if PACS1 p.R203W selectively induces protein pathway dysregulation and determine how these alterations compare to PACS1 loss of function.

### PACS1 p.R203W induces proteomic and kinase profiles distinct from PACS1 loss of function

We next performed unbiased quantitative proteomics analysis on day 29 iNs to identify molecular pathways altered by the p.R203W variant and compare them with PACS1 deficiency. We identified a total of 5,664 proteins with unique UniProtIDs across all genotypes. Principal component analysis on imputed and batch corrected matrices showed modest separation of PACS1^(+/R203W)^ samples from PACS1^(+/+)^ and PACS1^(-/-)^ samples (Figure S1A), supporting the idea that the pathogenic variant produces a molecular profile distinct from PACS1 loss of function.

To determine if PACS1 p.R203W induces proteomic changes distinct from PACS1 deficiency, we compared differentially abundant proteins (DAPs) across genotypes. We identified 228 DAPs in PACS1^(+/R203W)^ vs PACS1^(+/+)^, 197 DAPs in PACS1^(-/-)^ vs PACS1^(+/+)^, and 230 DAPs in PACS1^(-/-)^ vs PACS1^(+/R203W)^ iNs (Figure 2A, Table S1). While we observed some overlap, each comparison contained a comparable amount of unique DAPs, highlighting both convergent and divergent molecular consequences of the presence of the PACS1 pathogenic variant and the complete loss of PACS1 (Figure 2B). Among these, WDR37, a known PACS1 interactor was selectively decreased in PACS1^(-/-)^ iNs relative to PACS1^(+/+)^ and PACS1^(+/R203W)^ samples, which was confirmed by Western blot (Figure S1B-C).

**Figure 2.**
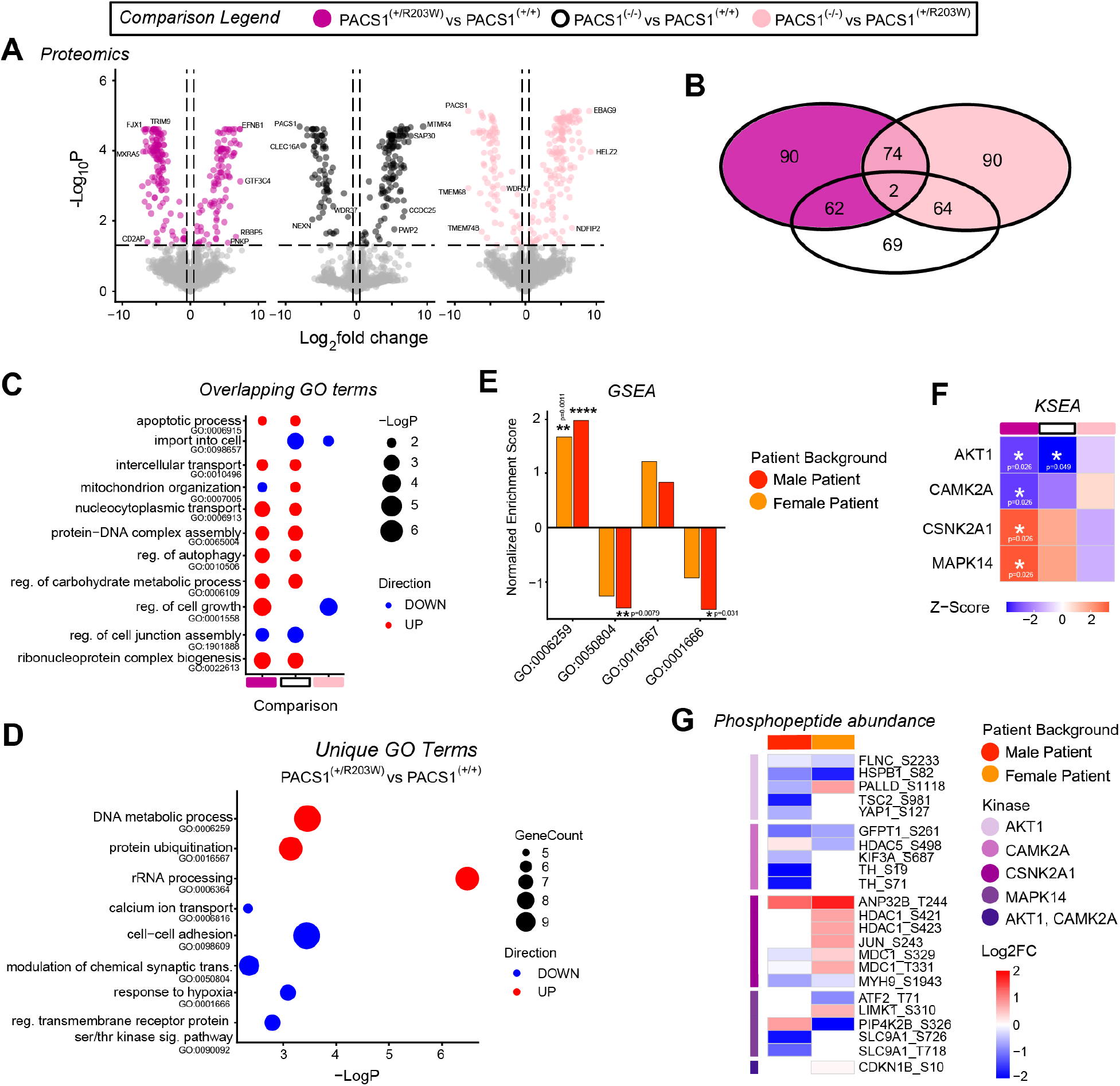
PACS1 p.R203W induces a proteomic and kinase signaling profile distinct from PACS1 loss of function. **(A)** Volcano plot of statistically significant DAPs with an absolute Log2FC > 0.5 across pairwise genotype comparisons (p-values≤ 0.05; n= 5372, 5443, 5429 proteins for PACS1^(+/R203W)^ vs PACS1^(+/+)^, PACS1^(-/-)^ vs PACS1^(+/+)^, PACS1^(-/-)^ vs PACS1^(+/R203W)^ comparisons, respectively). **(B)** Venn diagram showing the overlap of DAPs across pairwise genotype comparisons. **(C)** Dot plot showing statistically significant biological processes shared between the three pairwise comparisons; size of dot increases with increased statistical significance; red and blue indicate increase and decrease of each biological process in genotype being compared to another genotype. **(D)** Dot plot showing statistically significant biological processes unique to the PACS1^(+/R203W)^ vs PACS1^(+/+)^ comparison; statistical significance increases from left to right on X axis; size of dot is proportional to number of proteins identified within each biological process; red and blue indicate biological process is increased and decreased, respectively, in PACS1^(+/R203W)^ samples compared to PACS1^(+/+)^ samples. **(E)** Gene set enrichment analysis of male and female patient-derived iNs using Log2FC relative to their isogenic controls; Normalized Enrichment Scores for GO terms unique to PACS1^(+/R203W)^ vs PACS1^(+/+)^ comparison that follow predicted trends in patient-derived samples (significant p-values displayed, *p ≤ 0.05, **p ≤ 0.01, ****p ≤ 0.0001). **(F)** Kinase substrate enrichment analysis identifying predicted changes in kinase activity in across pairwise genotype comparisons; directionality of inferred kinase activity displayed by Z-score (significant p-values displayed, *p ≤ 0.05). **(G)** Validation of predicted kinase activity changes in male and female patient-derived iNs using Log2FC relative to their isogenic controls for targets downstream of AKT1, CAMK2A, CSNK2A1, and MAP14.

We performed Gene Ontology (GO) to identify biological processes enriched and diminished in each pairwise comparison. Interestingly, we found several overlapping GO terms in PACS1^(+/R203W)^ and PACS1^(-/-)^ samples relative to PACS1^(+/+)^ control. GO terms related to apoptosis (GO:0006915), intercellular transport (GO:0010496), nucleocytoplasmic transport (GO:0006913), protein-DNA complex assembly (GO:0065004) were enriched in PACS1^(+/R203W)^ and PACS1^(-/-)^, to name a few. A point of divergence was observed for mitochondrion organization (GO:0007005), which is diminished in PACS1^(+/R203W)^ iNs but enriched in PACS1^(-/-)^ iNs. Lastly, PACS1^(-/-)^ iNs were diminished in the biological process concerning import into the cell (GO:0098657) in comparison to both PACS1^(+/+)^ and PACS1^(+/R203W)^ iNs, highlighting a key aspect of PACS1 function that is conserved by the p.R203W variant and affected by loss of PACS1 function.

Next, we examined biological processes that were unique to each pairwise comparison (Table S2). When looking at GO terms enriched in PACS1^(+/R203W)^ relative to PACS1^(+/+)^, we saw an increase in DNA metabolic process (GO:0006259), protein ubiquitination (GO:0016567), and rRNA processing (GO:0006364), and decrease in calcium ion transport (GO:0006816), cell-cell adhesion (GO:0098609), modulation of chemical synaptic transmission (GO:0050804), response to hypoxia (GO:0001666), and regulation of transmembrane receptor proteins involved in Serine/Threonine kinase signaling pathways (GO:0090092) (Figure 2D). Altogether, the diversity of the biological processes affected by the p.R203W variant demonstrate the overall disruption to neuronal homeostasis that is occurring in our model for PACS1 syndrome. To validate these findings, we performed a targeted gene-set enrichment analysis (GSEA) on patient-derived iNs to determine if these highlighted biological processes unique to the PACS1^(+/R203W)^ vs PACS1^(+/+)^ comparison follow the expected trend when compared to their isogenic controls (Figure 2E). We observed that 4 of the 8 biological processes (GO:0006259, 0050804, 0016567 and 0001666) display the same trends in enrichment, consistent with our predicted alterations.

To determine whether these genotype specific differences in biological processes were accompanied by altered kinase signaling, we next performed phosphopeptide-enriched proteomic profiling. We identified a total of 9,336 unique phosphopeptides across all genotypes. Principal component analysis on imputed and batch corrected matrices showed modest separation of samples by genotype (Figure S1D). Pairwise comparisons identified 678 differentially abundant phosphopeptides (DAPhs) in PACS1^(+/R203W)^ vs PACS1^(+/+)^, 604 DAPhs in PACS1^(-/-)^ vs PACS1^(+/+)^, and 727 DAPhs in PACS1^(-/-)^ vs PACS1^(+/R203W)^ (Figure S1E). Overlap analysis revealed comparable amounts of unique DAPhs and associated proteins across comparisons, highlighting convergent as well as divergent effects of p.R203W variant and PACS1 deficiency on neuronal signaling pathways (Figure S1F).

To identify kinases upstream of our observed phosphorylation changes, we performed kinase-substrate enrichment analysis (KSEA) (38). This analysis predicted decreased activity of AKT1 and CAMK2A, and increased CSNK2A1 and MAPK14 activity in PACS1^(+/R203W)^ iNs relative to PACS1^(+/+)^ controls (Figure 2F). Interestingly, decreased AKT1 activity is also predicted in PACS1^(-/-)^ iNs, suggesting a potential point of convergence between the p.R203W variant and PACS1 deficiency. To validate these findings, we examined the abundance of downstream kinase substrates in patient-derived iNs and observed changes consistent with the predicted alterations of AKT1, CAMK2A, CSNK1A and MAPK14 activity relative to their isogenic controls (Figure 2G). Importantly, none of these kinases were DAPs in our global proteomic dataset (Figure S1G), indicating that the observed signaling changes likely reflect altered kinase activity rather than differences in protein expression. Collectively, the global proteomics and KSEA analyses reveal a molecular signature unique to PACS1^(+/R203W)^ iNs that is distinct from PACS1 deficiency. These findings support a model in which the PACS1 p.R203W variant alters signaling networks through mechanisms beyond simple loss of PACS1 function, consistent with a non-loss-of-function pathogenic mechanism.

### PACS1^(+/R203W)^ converges with PACS1^(-/-)^ on synaptic phenotypes but induces distinct morphological changes

Since our GO analysis identified a decreased enrichment of biological processes related to synaptic transmission in our PACS1^(+/R203W)^ iNs relative to PACS1^(+/+)^ iNs, we quantified synaptic puncta and used immunofluorescence to look at markers for neuronal maturation. Interestingly, both PACS1^(+/R203W)^ and PACS1^(-/-)^ iNs have decreased number of PSD95 puncta and colocalizing PSD95/Synapsin1 puncta relative to PACS1^(+/+)^ controls, while fewer Synapsin1 puncta was observed only in PACS1^(+/R203W)^ iNs (Figure 3A-B). Consistent with these findings, both PACS1^(+/R203W)^ and PACS1^(-/-)^ iNs have decreased MAP2 and NeuN fluorescence intensity, suggesting delayed or impaired neuronal maturation compared to PACS1^(+/+)^ iNs (Figure 3C-E) (39–43). Despite these changes, gross morphology was largely preserved. No difference in branching complexity, total neurite length, or maximum neurite length (Figure 3F-H) was observed. Interestingly, PACS1^(+/R203W)^ iNs showed a unique reduction in soma area that was not observed in PACS1^(-/-)^ iNs (Figure 3I). While PACS1^(+/R203W)^ and PACS1^(-/-)^ iNs share defects in synaptic density and possibly have impaired maturation timelines, the p.R203W variant also induces a unique morphological phenotype. Altogether, these findings indicate that p.R203W induces a combination of overlapping and divergent phenotypes relative to PACS1 deficiency, further supporting that p.R203W does not operate via a simple LoF mechanism.

**Figure 3.**
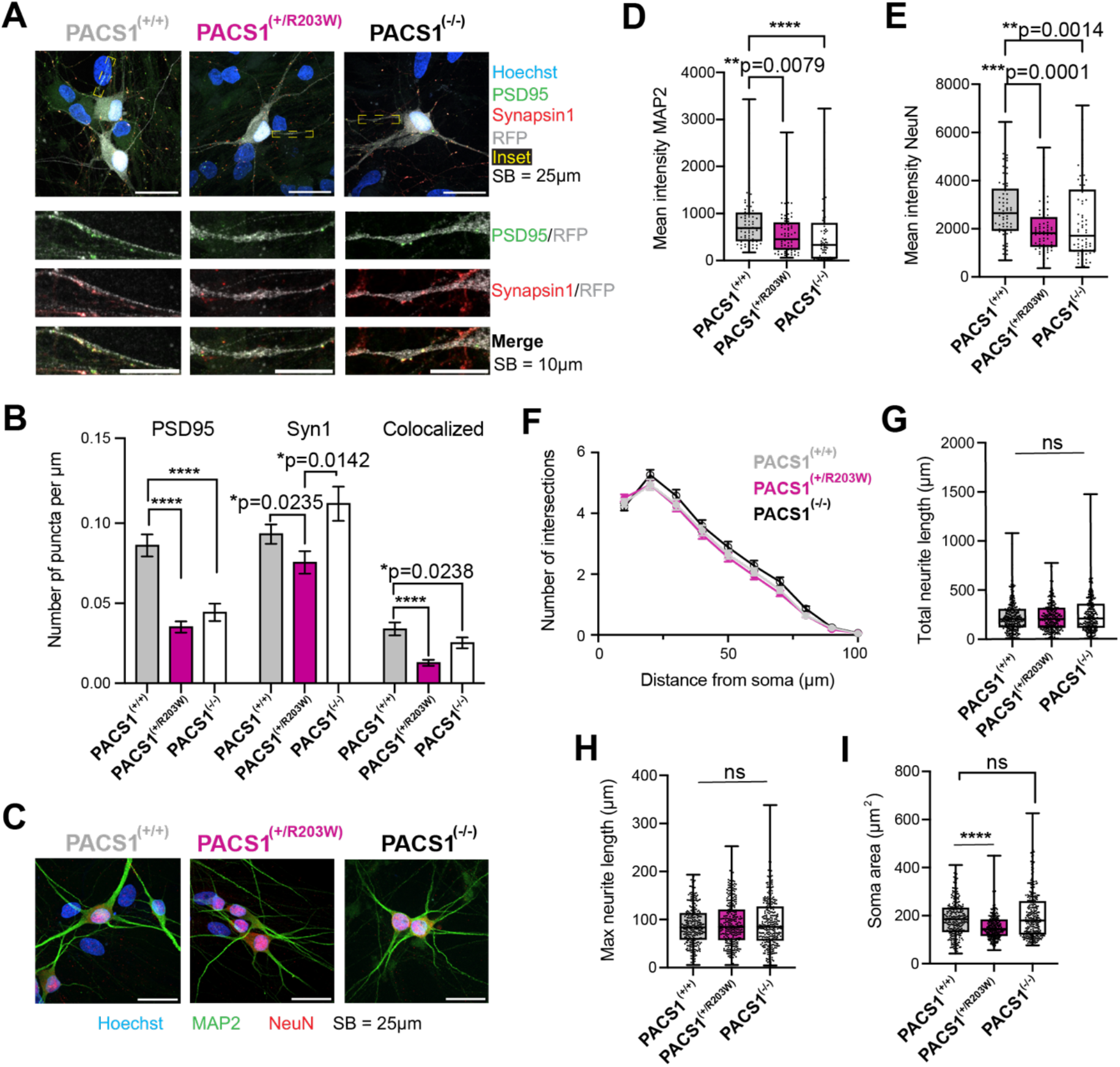
PACS1^(+/R203W)^ converges with PACS1^(-/-)^ on synaptic phenotypes but induces distinct morphological changes. **(A)** Representative images of PSD95 and Synapsin1 puncta in sparsely labeled PACS1^(+/+)^, PACS1^(+/R203W)^ and PACS1^(-/-)^ d35 iNs; SB = 25μm, inset 10μm. **(B)** Quantification of PSD95 puncta (Kruskal-Wallis with Dunn’s multiple comparisons; p_Kruskal_<0.0001; p_R203W_<0.0001, p_KO_<0.0001; n=70 cells per genotype), Synapsin1 puncta (Kruskal-Wallis with Dunn’s multiple comparisons; p_Kruskal_=0.0066; p_R203W_=0.0235, p_KO_>0.9999; n=70 cells per genotype) and colocalizing puncta (Kruskal-Wallis with Dunn’s multiple comparisons; p_Kruskal_<0.0001; p_R203W_<0.0001, p_KO_=0.0581; n=70 cells per genotype), normalized to total neurite length, across the three genotypes. **(C)** Representative images of MAP2 and NeuN in PACS1^(+/+)^, PACS1^(+/R203W)^ and PACS1^(-/-)^ d35 iNs, SB = 25μm. **(D)** Quantification of mean fluorescence intensity of MAP2 across genotypes (Kruskal-Wallis with Dunn’s multiple comparisons; p_Kruskal_<0.0001; p_R203W_=0.0079, p_KO_<0.0001; n=70 cells per genotype). **(E)** Quantification of mean fluorescence intensity of nuclear NeuN across all genotypes (Kruskal-Wallis with Dunn’s multiple comparisons; p_Kruskal_<0.0001; p_R203W_=0.0001, p_KO_=0.0014; n=70 cells per genotype). **(F)** Sholl analysis measuring the branching complexity of d35 iNs from all three genotypes (2way ANOVA with Tukey’s multiple comparisons test; all p-values > 0.05; n=280 cells per genotype. **(G)** Quantification of total neurite length (Kruskal-Wallis with Dunn’s multiple comparisons; p_Kruskal_=0.6601; p_R203W_>0.9999, p_KO_>0.9999; n=280 cells per genotype), **(H)** maximum neurite length (Kruskal-Wallis with Dunn’s multiple comparisons; p_Kruskal_=0.6882; p_R203W_>0.9999, p_KO_>0.9999; n=280 cells per genotype), and **(I)** soma size (Kruskal-Wallis with Dunn’s multiple comparisons; p_Kruskal_<0.0001; p_R203W_<0.0001; p_KO_>0.9999; n=280 cells per genotype) of d35 iNs from all three genotypes.

### PACS1 p.R203W results in stimulus dependent shift in excitability curve

We performed whole cell patch-clamp recordings on day 35-42 iNs to determine how the p.R203W variant affects neuronal function and how these defects compare to PACS1 deficiency (Figure 4A). Using a current clamp step protocol, we observed a shift in the excitability curve phenotype in PACS1^(+/R203W)^ iNs, characterized by significantly increased action potential firing frequencies at higher current injections (150-170 pA) relative to PACS1^(+/+)^ iNs (Figure 4B-C). In contrast, PACS1^(-/-)^ iNs exhibited firing properties comparable to PACS1^(+/+)^ iNs across all stimulus ranges, despite the molecular and synaptic alterations described above. These findings indicate that the observed shift in excitability is directly associated with the expression of the p.R203W variant and is not recapitulated by loss of PACS1, further supporting a non-loss-of-function mechanism for PACS1 syndrome pathogenesis.

**Figure 4.**
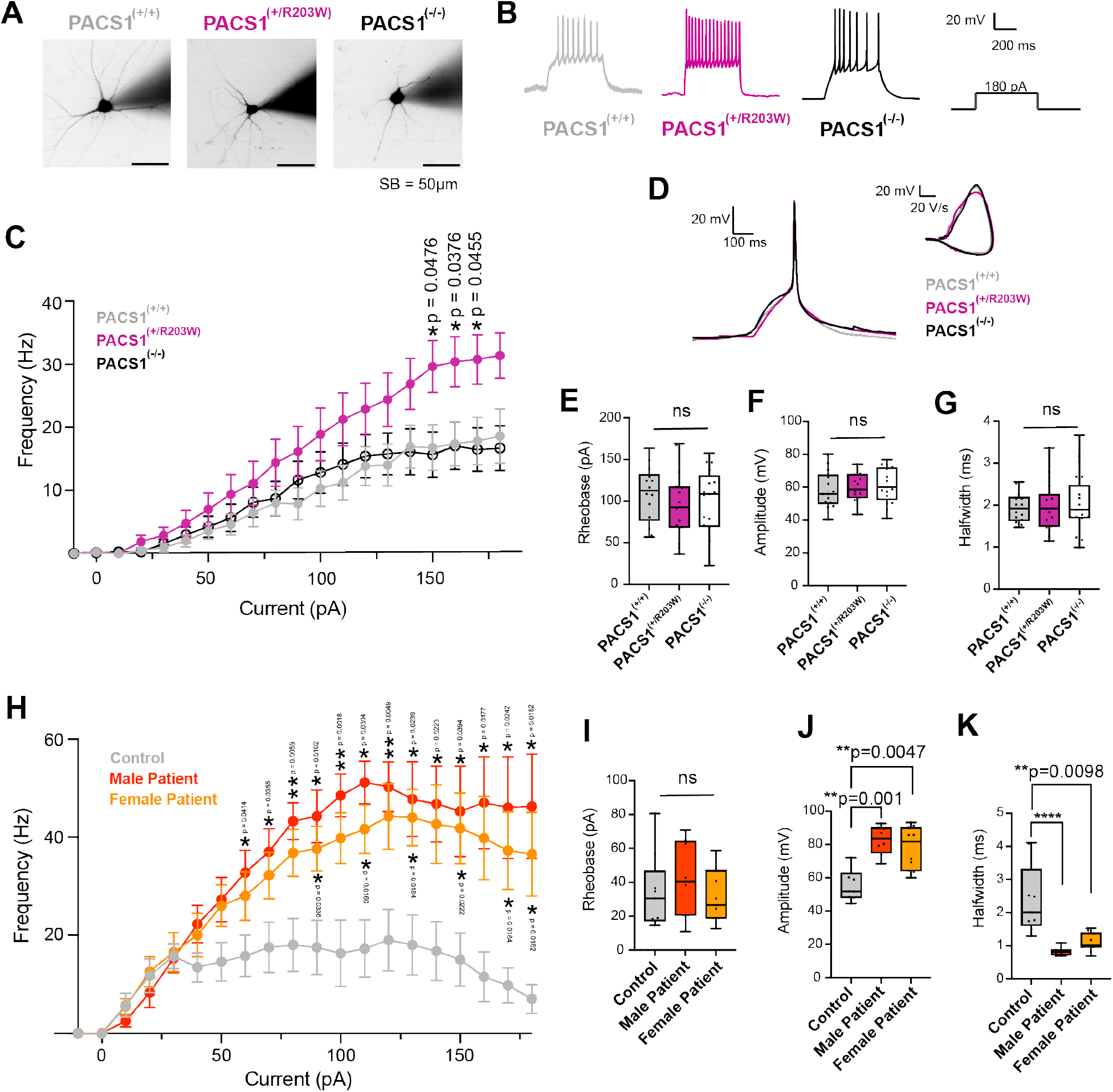
PACS1 p.R203W results in a stimulus dependent shift in excitability curve. **(A)** Representative images of patched PACS1^(+/+)^, PACS1^(+/R203W)^ and PACS1^(-/-)^ d35-42 iNs loaded with Alexa-Fluor488; SB = 50μm. **(B)** Representative traces of action potential spiking frequency at 180pA across the three genotypes. **(C)** Excitability curve across the three genotypes (2way ANOVA with Tukey’s multiple comparisons test; significant p-values displayed; n= 13, 16, 17 for PACS1^(+/+)^, PACS1^(+/R203W)^ and PACS1^(-/-)^, respectively. **(D)** Representative traces of action potentials and phase plane plots across the three genotypes, highlighting minimal variation. **(E)** Quantification of rheobase (Kruskal-Wallis with Dunn’s multiple comparisons; p_Kruskal_=0.6264 p_R203W_=0.6822, p_KO_>0.9999; n= 14, 15, 17 for PACS1^(+/+)^, PACS1^(+/R203W)^ and PACS1^(-/-)^, respectively), **(F)** amplitude (Kruskal-Wallis with Dunn’s multiple comparisons; p_Kruskal_=0.6658; p_R203W_>0.9999, p_KO_=0.7480; n= 14, 15, 17 for PACS1^(+/+)^, PACS1^(+/R203W)^ and PACS1^(-/-)^, respectively) and **(G)** halfwidth (Kruskal-Wallis with Dunn’s multiple comparisons; p_Kruskal_=0.9871; p_R203W_>0.9999, p_KO_>0.9999; n= 14, 15, 17 for PACS1^(+/+)^, PACS1^(+/R203W)^ and PACS1^(-/-)^, respectively) corresponding to action potentials from PACS1^(+/+)^, PACS1^(+/R203W)^ and PACS1^(-/-)^ iNs. **(H)** Current clamp step protocol plot of action potential spiking frequency against current injection across the three genotypes (2way ANOVA with Tukey’s multiple comparisons test; significant p-values displayed, *p ≤ 0.05, **p ≤ 0.01; n= 8, 8, 11 for control, male patient, and female patient, respectively. **(I)** Quantification of rheobase (Kruskal-Wallis with Dunn’s multiple comparisons; p_Kruskal_=0.5331; p_male_=0.6396, p_female_>0.9999; n= 9, 8, 11 for control, male patient, and female patient, respectively respectively), **(J)** amplitude (Kruskal-Wallis with Dunn’s multiple comparisons; p_Kruskal_=0.0008; p_male_=0.0010, p_female_=0.0047; n= 9, 8, 11 for control, male patient, and female patient, respectively respectively) and **(K)** halfwidth (Kruskal-Wallis with Dunn’s multiple comparisons; p_Kruskal_<0.0001; p_male_<0.0001, p_female_=0.0098; n= 9, 8, 11 for control, male patient, and female patient, respectively) corresponding to action potentials from summary control, male patient and female patient-derived iNs. All current-clamp and voltage-clamp quantifications are summarized in Tables S3 and S4.

To identify mechanisms underlying the PACS1^(+/R203W)^ excitability phenotype, we next examined action potentials properties using a minimal stimulation protocol (Figure 4D). We observed no difference in threshold, rheobase, amplitude, halfwidth and afterhyperpolarization across genotypes (Figures 4E-G, Table S3), suggesting that the increased firing rates observed in PACS1^(+/R203W)^ iNs cannot explained by alterations in these intrinsic membrane properties. We then used patient-derived iNs (PACS1^(+/R203W)^) and performed excitability curves, which phenocopied the observed leftward excitation shift seen in the PACS1^(+/R203W)^ knock-in line, but exhibited an exaggerated phenotype, with higher firing frequencies occurring at lower current injections (Figure 4H). Unlike our knock-in line, patient-derived iNs had significantly increased action potential amplitudes and decreased halfwidths, while the threshold and rheobase remained unchanged (Figure 4I-K, Table S4). These findings suggest that genetic background may modulate the severity of PACS1-associated neuronal dysfunction.

Lastly, to determine whether sodium and potassium channels contributed to the observed shift in excitability, we performed voltage-clamp recordings. No differences in capacitance-adjusted Max Na^+^ and Max K^+^ currents, input resistance or capacitance were observed among PACS1^(+/+)^, PACS1^(+/R203W)^ and PACS1^(-/-)^ isogenic iNs (Table S3). Similarly, patient-derived iNs were comparable to the control except for a slightly increased capacitance in the Male Patient line (Table S4). Overall, these findings suggest that the altered excitability of PACS1^(+/R203W)^ iNs is not driven by differences in membrane properties or ion-channel presence and/or function. Instead, the enhanced excitability phenotype in patient-derived iNs raises the possibility that genetic modifiers could influence disease severity in PACS1 syndrome.

### PACS1 p.R203W preserves core interactions while remodeling the PACS1 interactome

Our global and phosphopeptide-enriched proteomic analyses revealed that the pathogenic PACS1 p.R203W variant induces a unique molecular landscape distinct from PACS1 deficiency, which results in a stimulus dependent shift in neuronal excitability. This raises the question of how the p.R203W variant alters PACS1 function. Structural modeling using AlphaFold3 predicted that residue 203 undergoes a marked positional shift, changing from a surface exposed Arginine in PACS1(R203) to a Tryptophan located near the hydrophobic core in PACS1(W203) (Figures 5A, S2A). This observation suggested that this predicted structural change in PACS1(W203) could also alter PACS1 protein interactions. To test this possibility, we performed an *in silico* coimmunoprecipitation (coIP) using AlphaPulldown with our iN proteome as prey (Figure 5B) (44). After filtering proteins predicted to interact with GFP, we identified 142 candidate interactors for PACS1(R203)-GFP (Figures 5C, S2B, Table S5). GO analysis revealed enrichment for pathways involved in RNA and nucleotide metabolism, protein synthesis and ribosome biology, protein processing and modification, organelle and cellular organization, energy and carbohydrate metabolism, autophagy and cellular degradation, signaling and apoptosis, and lipid and steroid metabolism and signaling (Figure 5D). Notably, 28 were associated with NDDs and 11 were Autism Spectrum Disorder (ASD) risk genes (Figure 5E). Interestingly, the biological processes associated with candidate interactors of PACS1(R203)-GFP were related to non-specific cellular biology, even in the context of the neuronal proteome.

**Figure 5.**
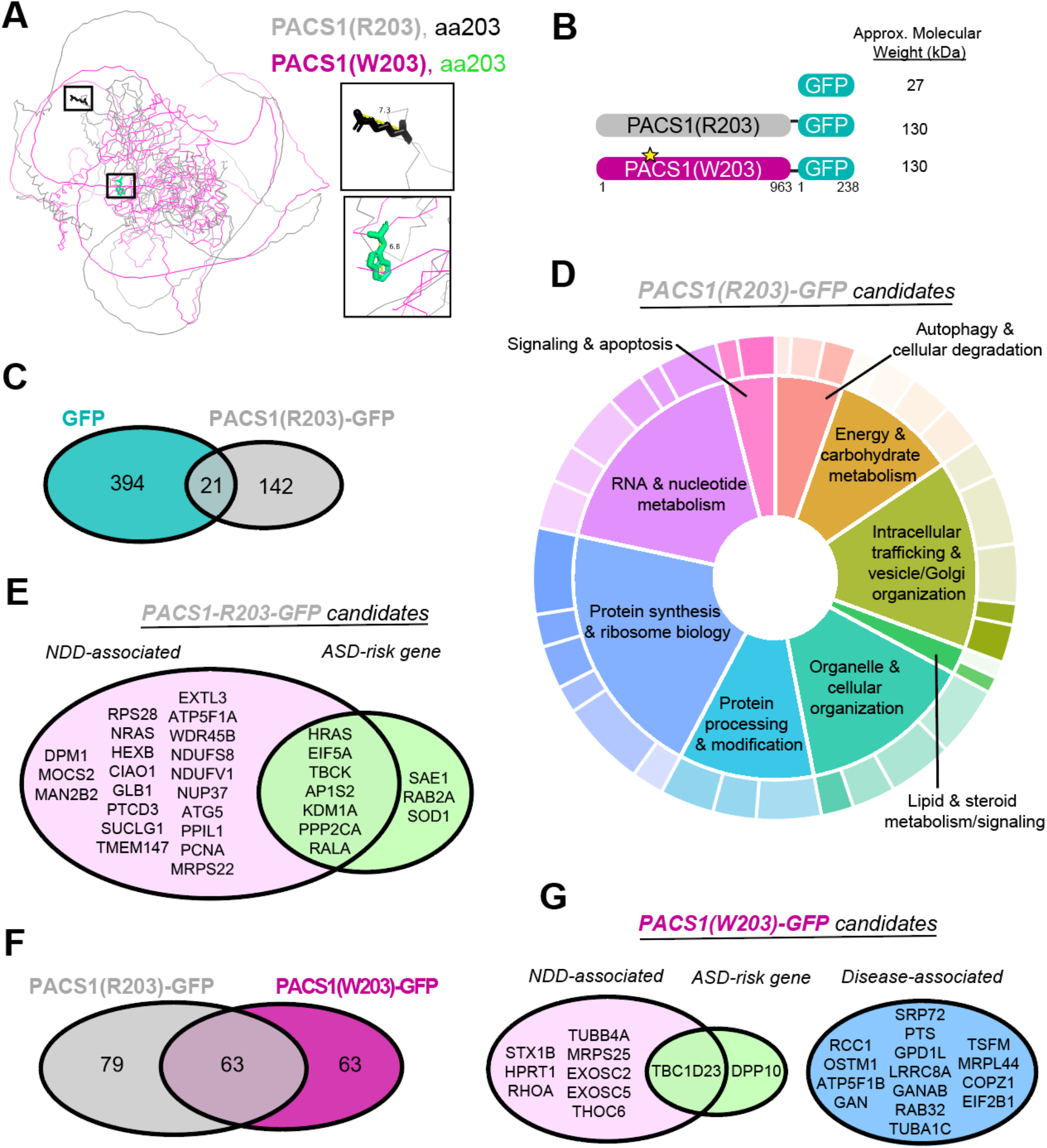
PACS1 p.R203W is predicted to alter interactions while preserving core PACS1 interactome. **(A)** AlphaFold3 structural models of PACS1 containing either R203 of W203 residue, highlighting predicted structural changes and local environment of residue 203. **(B)** Schematic of GFP-tagged constructs used to analyze the PACS1 interactome. **(C)** Overlap of predicted candidate interactors for GFP and PACS1(R203)-GFP. **(D)** Gene ontology analysis of predicted PACS1(R203)-GFP candidate interactors, grouped by shared molecular function; sizing of pie slice dependent on gene count for that GO-term/group. **(E)** PACS(R203)-GFP candidate interactors associated with NDDs and ASD. **(F)** Overlap of predicted candidate interactors for PACS1(R203)-GFP and PACS1(W203)-GFP. **(H)** PACS1(W203)-GFP specific candidate interactors associated with diseases. All data shown are representative of results from *in silico* coimmunoprecipitation. All predicted candidate interactors listed in Table S5.

We next validated these findings experimentally using coIP followed by mass spectrometry in HEK293T cells overexpressing these GFP-tagged constructs. Successful pulldown was confirmed by Western blotting for PACS1, GFP, and the known PACS1 interactor WDR37, which was detected in the bound fraction isolated from both PACS1(R203)-GFP and PACS1(W203)-GFP samples (Figure S2C) (29, 30). After filtering proteins detected in GFP-only control, 88 candidate interactors associated with PACS1(R203)-GFP were identified (Figure S2D, Table S6). Consistent with the *in silico* analysis, GO enrichment of experimentally identified interactors revealed pathways involved in RNA metabolism, protein homeostasis, protein localization, stress responses, autophagy, and chromatin organization (Figure S2E).

Comparison of the *in silico* candidate interactors PACS1(R203)-GFP and PACS1(W203)-GFP revealed substantial overlap with 63 shared candidates. However, 79 proteins were unique to PACS1(R203)-GFP and 63 remained unique to PACS1(W203)-GFP (Figure 5F, Table S5). Several of the PACS1(W203)-GFP candidate interactors have reported disease associations, including NDDs and ASD (Figure 5G), suggesting that the variant may acquire novel interactions that contribute to pathogenesis.

Altogether these findings suggest that the PACS1 p.R203W pathogenic variant does not abolish the core of the PACS1 interactome, as there remains a large degree of overlap between candidate interactors identified for PACS1(R203)-GFP and PACS1(W203)-GFP. Instead, the pathogenic variant appears to selectively remodel the PACS1 interactome through the acquisition and loss of specific interactors. These altered interactions may contribute to the distinct molecular and functional phenotypes observed in PACS1^(+/R203W)^ iNs and provide a potential mechanistic basis to PACS1 syndrome pathogenesis.

## DISCUSSION

PACS1 syndrome is caused by the recurrent *de novo* pathogenic variant p.R203W, yet the molecular mechanism by which this single amino acid substitution disrupts neurodevelopment remains incompletely understood. While previous studies have suggested that the PACS1 p.R203W pathogenic variant operates by a GoF mechanism, direct comparison to PACS1 knockdown and/or knockout models have been lacking. Our findings demonstrate that there is a level of phenotypic overlap between PACS1^(+/R203W)^ and PACS1^(-/-)^ iNs, yet the p.R203W variant induces distinct molecular phenotypes that result in a unique functional phenotype that is not recapitulated in PACS1^(-/-)^ iNs. Together our results provide the first evidence that PACS1 syndrome is not caused by a simple loss of PACS1 function mechanism and instead supports a non-loss-of-function mechanism. This non-loss-of-function mechanism remains to be determined.

Distinguishing the effects of p.R203W from PACS1 deficiency is an important step in identifying therapeutic options. Recent N-of-1 trials using non-allele-specific ASOs have highlighted the need to better define the consequences of reducing PACS1 abundance. Our data indicate that while there is a degree of phenotypic convergence, PACS1 deficiency does not functionally phenocopy the PACS1 p.R203W pathogenic variant, suggesting that PACS1^(+/R203W)^ and PACS1^(-/-)^ iNs operate via divergent mechanisms. Moreover, a better understanding of the effects of p.R203W on PACS1 function remains an important part of informing and developing therapeutic targets for PACS1 syndrome patients.

Given that PACS1 is described for its canonical function as a tGN sorting protein and that Golgi architecture phenotypes have been previously described (22, 31), we initially anticipated identifying defects in Golgi organization and/or endosomal trafficking. To our surprise, we did not observe any difference in Golgi structure or positioning, distribution of EEA1-positive vesicles or PACS1 localization between PACS1^(+/+)^ and PACS1^(+/R203W)^ iNs. These findings suggest that major alterations in canonical PACS1 trafficking functions are not a prominent feature of our human neuronal model and may be context dependent (45).

More broadly, our results support the growing view that PACS1 functions beyond its canonical role in intracellular trafficking. Since its original description as a tGN sorting protein, PACS1 has been implicated in diverse cellular processes including nuclear transport, genomic stability, calcium regulation in the endoplasmic reticulum, regulation of microtubule dynamics, to name a few (22, 23, 29, 34). Consistent with this expanded view, our findings highlight candidate interactors that have never been reported in the PACS1 literature, including proteins associated with autophagy, proteostasis, and RNA metabolism. Many of these proteins have established links to NDDs, suggesting that PACS1 may participate in broader cellular networks relevant to brain development and disease.

At the molecular level, PACS1^(+/R203W)^ iNs exhibited a unique global and phosphopeptide-enriched proteomic signature characterized by a broad dysregulation of protein pathways and altered kinase signaling. These changes were accompanied by reduced synaptic density and stimulus dependent increase in neuronal excitability, a phenotype that was absent in PACS1^(-/-)^ iNs. The electrophysiology findings corroborate that PACS1^(+/R203W)^ iNs are in a state of dysregulation, as at higher current injections cells begin to fire action potentials erratically. Importantly, patient-derived neurons recapitulated and, in some cases, amplified these functional abnormalities, suggesting that genetic background may influence disease severity. Although the precise mechanisms underlying altered neuronal excitability remain unclear, the convergence of altered signaling pathways, synaptic abnormalities, and electrophysiological dysfunction points to widespread disruption of neuronal homeostasis in PACS1 syndrome.

Lastly, our *in silico* coIP, and subsequent experimental validation provide a potential mechanistic explanation for the distinct molecular and functional phenotypes induced by p.R203W. Predicted AlphaFold3 structures for PACS1(R203) and PACS1(W203) suggested conformational changes and positioning of residue 203, bringing to question the potential impact of the pathogenic variant on overall PACS1 structure and function. While several predicted candidate interactors were shared between PACS1(R203)-GFP and PACS1(W203)-GFP, both constructs had unique candidate interactors. These findings suggest that p.R203W may preserve aspects of canonical PACS1 function while selectively remodeling its interactome through the acquisition and/or loss of specific binding partners. Such interactors could contribute to the distinct molecular signature and altered excitability observed in PACS1^(+/R203W)^ iNs. Although these candidate interactors require further validation, they provide a novel framework for future mechanistic studies and may reveal new opportunities for therapeutic intervention.

In summary, our study demonstrates that the PACS1 syndrome causative variant p.R203W does not act through a simple LoF mechanism. Instead, the pathogenic variant produces a unique molecular and functional state characterized by broad protein pathway dysregulation and altered kinase signaling, ultimately resulting in a stimulus dependent shift in neuronal excitability. These findings refine our understanding of PACS1 syndrome pathogenesis and establish a foundation for future studies aimed at identifying disease relevant pathways and targeted therapies to treat PACS1 syndrome patients.

## METHODS

### Sex as a biological variable

Both female and male iPSC lines were included in this study. The primary comparison of PACS1^(+/+)^, PACS1^(+/R203W)^ and PACS1^(-/-)^ genotypes used isogenic female iPSC lines to minimize the variability of genetic background. Findings were validated in iNs derived from a male and female affected by PACS1 syndrome in comparison to their CRISPR Cas9 generated isogenic controls. PACS1 syndrome affects both sexes, and there is currently no evidence that disease severity is influenced by sex. Sex was not evaluated as an independent variable.

### Cell culture

#### iPSC culture

Three PACS1^(+/+)^, three PACS1^(+/R203W)^ and three PACS1^(-/-)^ iPSC lines were used in these experiments. All PACS1^(+/+)^ and PACS1^(+/R203W)^ have been previously described (35). Briefly, one PACS1^(+/+)^ line was derived from a female control (GM03651; Coriell Institute for Medical Research) and the corresponding PACS1^(+/R203W)^ was generated by introducing the p.R203W variant using CRISPR Cas9. The remaining two PACS1^(+/R203W)^ lines were derived from a male and female with PACS1 syndrome (GM27159 & GM27161; WiCell Research Institute), and the p.R203W variant was corrected using CRISPR Cas9 to create isogenic controls in the patient genetic background. All three PACS1^(-/-)^ lines were generated using the previously described CRISPR Cas9 approach, two of which were generated in the PACS1^(+/+)^, female control line, and the third line was generated in a male control line (PGP1 background; Synthego). iPSCs were maintained in mTeSR+ medium (Stem Cell Technologies, 05825) on plates coated with Matrigel (Corning, 354234) dissolved in 1x DMEM at 0.1 mg/mL. Cells were passaged as needed using ReLeSR (Stem Cell Technologies, 05782). Cell lines were regularly tested for mycoplasma contamination. Cell lines were authenticated using a combination of Sanger Sequencing for genotype confirmation, short tandem repeat (STR) profiling, in-Process Control System-digital Pluripotent Stem Cell (iCS-digital PSC, StemGenomics) for chromosomal abnormalities, and immunofluorescence for pluripotency markers.

#### iNeuron differentiation

iPSC lines were differentiated into a predominantly excitatory population of neurons through the overexpression of transcription factor Ngn2 in a doxycycline inducible lentiviral system, previously described (35, 46).

#### iNeuron culture

iNs were plated in plates coated with 0.2 mg/mL Matrigel. If coverslips were to be used, coverslips were initially acid washed in 1M HCl and sterilized by autoclave before being coated with 0.2 mg/mL Matrigel. If iNs were to be co-cultured with rat primary astrocytes (Thermo Scientific, N7745-100), astrocytes were plated in a monolayer 2-3 days prior to iN plating (maintained in 1xDMEM with 15% FBS until iNs plated). iNs were cultured in plating/maturation media with partial media changes on a M/W/F schedule. iNs intended for immunofluorescence were sparse-labeled using an AAV1-hSynapsin1-mScarlett (Addgene, 131001-AAV1) at an MOI 2500 at d25 of culture.

*HEK293T culture:* HEK293T (American Type Culture Collection, CRL-1573) were cultured in 1x DMEM with 10% FBS. Cells were passaged as needed using 0.05% Trypsin-EDTA. Mycoplasma testing was done routinely.

### Western blots

iNs (d29) were collected in 1x RIPA buffer (Cell Signaling Technology, 9806) supplemented with protease and phosphatase inhibitors (Thermo Scientific, 23227). Protein concentration was determined using the Pierce microBCA protein assay kit (Thermo Scientific, 23235), and lysates were denatured by adding 4x Laemmli buffer (Bio-Rad, 1610747) with βmercaptoethanol and boiling at 95°C. 10μg of protein extract was loaded into polyacrylamide gels (Bio-Rad, 4-15% or 7.5% gels) and transferred to PVDF membranes (Thermo Scientific, 88518). Prior to immunoblotting, all membranes were stained for total protein (LiCor, 926-11016). Membranes were probed with antibodies against PACS1 (Sigma Aldrich, HP1038914, 1:1000), WDR37 (Sigma Aldrich, HPA037376, 1:1000), acetylated tubulin (Sigma Aldrich, T7451, 1:1000), GAPDH (Cell Signaling Technology, 97166S, 1:1000), and GFP (Abcam, ab13970, 1:5000). Membranes were incubated with the following secondary antibodies, all at a dilution of 1:10,000: goat anti-rabbit IRDye 680RD (Li-Cor, 926-68071) goat anti-rabbit IRDye 800CW (Li-Cor, 926-32211), goat anti-mouse IRDye 680RD (Li-Cor, 926-68070) goat anti-mouse IRDye 800CW (Li-Cor, 926-32210). Immunoblots were visualized using the Odyssey Fc Imaging System (Li-Cor) and quantified using Image Studio (Li-Cor).

### Immunofluorescence

Samples were prepared for n=2 biological replicates for each genotype (PACS1^(+/+)^, PACS1^(+/R203W)^, PACS1^(-/-)^). iNs were co-cultured with rat primary astrocytes until d35. Samples were fixed in 4% paraformaldehyde, permeabilized with PBT (1x PBS with 0.3% Triton X-100), and blocked with 5% Normal Donkey Serum (SouthernBiotech, OB003001) diluted in PBT. Samples were incubated primary antibody overnight at 4°C. Primary antibodies used were PACS1(1:200), Giantin (Abcam, ab37266, 1:200), EEA1 (Abcam, ab70521, 1:200), MAP2 (Invitrogen, 13-1500, 1:400), NeuN (Thermo Scientific, PA5-78499, 1:200), PSD95 (NeuroMab, 75-028, 1:200), Synapsin (Cell Signaling Technology, 5297S, 1:200), and RFP (ProteinTech, 5F8, 1:1000). Secondary antibodies used were donkey anti-mouse488 (Invitrogen, A-21202), donkey anti-rat594 (Invitrogen, A-21209) and donkey anti rabbit 647 (Invitrogen, A-31573). Samples were stained with Hoechst 33342 (Invitrogen, H3570, diluted 1:3000) before mounting with Fluoromount-G (Thermo Scientific, 00-4958-02).

### Microscopy

Images were taken at 100x on a Nikon CSU-W1 spinning disk microscope in the Center for Advanced Microscopy at Northwestern University. Briefly, 35 RFP-positive cells were identified (per genotype, per coverslip) at 60x using the epifluorescence microscope. Each 4μm Z-stack was centered at the widest cross-section of the cell’s nucleus, with 0.2μm Z-steps.

### Image analysis

All images were blinded prior to analysis. Image analysis was performed using ilastik (v1.4.2) for segmentation and binary mask generation and FIJI (v1.54t) plugins and custom macros. For each stain, all images were split into DAPI, GFP, RFP, FarRed channels. Max projections were made for the RFP channel (sparse labeled cells); average projections were generated for remaining channels. Sparse-labeled neurons were manually traced to generate soma and neurite binary masks. Sholl analysis was performed using Simple Neurite Tracer using the soma centroid and neurite binary mask. Neurite masks were skeletonized using the Skeletonize plugin. Analyze Skeleton (2D/3D) was used to quantify the total neurite length and longest neurite path. Binary masks for nuclei, soma, neurites, whole cell, Golgi, and EEA1 vesicles were used to measure object area and the mean fluorescence intensity of PACS1, MAP2, and NeuN within those objects. Golgi perinuclear distance was quantified by measuring the distance between the nuclear centroid and Golgi centroid. Average projections of PSD95 and Synapsin1 were generated into binary masks using ilastik. Puncta were quantified within the neurite binary mask. Colocalized puncta were quantified by counting the number of overlapping PSD95 and Synapsin1 objects within the neurite binary mask. All puncta quantifications were normalized to total neurite length of the cell.

### Proteomics

Proteomics samples were collected for 3 biological replicates for isogenic lines generated in control female background (n=3 PACS1^(+/+)^, n=3 PACS1^(+/R203W)^, n=3 PACS1^(-/-)^). Samples for patient-derived iNs were also collected with their appropriate isogenic control (n=1 female patient, n=1 female isogenic control, n=1 male patient, n=1 male isogenic control). iNs (d29) were collected in urea buffer (8M urea, 100mM Tris-HCl, pH 8) supplemented with protease and phosphatase inhibitors. Samples were sonicated on ice with 20% amplitude for two 15 sec pulses. Lysates were cleared and quantified. 500μg of protein for each sample was used for mass spectrometry preparation. Reduction, alkylation, enzymatic digestion, and peptide cleanup were performed using adaptations to a previously described workflow (47).

#### Global proteomics samples

An aliquot of each desalted peptide digest was reserved for global proteomic analysis before phosphopeptide enrichment. Assuming 80% peptide recovery from 500µg of starting protein, approximately 400µg of peptide was expected per sample. Dried peptide digests were reconstituted in 50µL of solvent A (0.1% formic acid). A 5µL aliquot was diluted with 45µL of solvent A to generate a tenfold dilution. A 2µL aliquot of each diluted sample was injected for LC-MS/MS analysis, corresponding to an estimated peptide load of approximately 1.6 µg per run.

#### Phosphopeptide-enriched proteomics samples

Phosphopeptides were enriched using the Pierce High-Select TiO₂ Phosphopeptide Enrichment Kit (Thermo Scientific, A32993) according to the manufacturer’s protocol. Enriched phosphopeptides were dried by vacuum centrifugation and stored at -80°C until analysis. Before LC-MS/MS analysis, sample were reconstituted in 10µL of solvent A, and 2µL was injected per run. The phosphopeptide amount was not measured directly; the anticipated yield was estimated to be 1% of the starting peptide material.

### HEK293T coimmunoprecipitation

HEK293T cells were infected with lentiviruses for GFP, R203-GFP and W203-GFP (lentiviral constructs purchased from Vector Builder, VB240318-1512hgx, VB240318-1502cpx, VB240318-1520uuy; lentivirus generated in Northwestern University GET-iN Core). Cells were collected and lysed, and GFP-tagged proteins were co-immunoprecipitated using ChromoTek GFP-Trap Magnetic Agarose beads (ProteinTech, gtma-20) following the manufacturer’s standard immunoprecipitation protocol. Proteins captured by the beads went through on-bead digestion based on manufacturer’s standard protocol which is a modification of the workflow previously described (48). Peptides were dried by vacuum centrifugation and desalted using ZipTips according to the Northwestern Proteomics ZipTip protocol and dried again. For LC-MS/MS analysis, peptides were reconstituted in 20µL mobile phase A (5% acetonitrile, 0.1% formic acid), mixed, sonicated, and centrifuged; a 5µL aliquot was injected per analysis.

### Mass spectrometry

#### Proteomics

Global and phosphopeptide-enriched samples were analyzed using a Vanquish Neo nanoflow liquid chromatography (LC) system coupled to an Orbitrap Exploris 480 mass spectrometer. Samples were loaded using a trap-and-elute configuration with a 300-µm internal diameter × 5-mm trapping column and separated on a 75-µm internal diameter × 25-cm analytical column. The analytical flow rate was maintained at 300 nL/min. The autosampler temperature was maintained at 7°C, and trap loading and washing were performed at 50 µL/min. Mobile phase A was 0.1% formic acid and mobile phase B was 80% acetonitrile containing 0.1% formic acid. Samples were reconstituted for injection in 5% acetonitrile containing 0.1% formic acid. The 120-min LC method began at 0% B, increased to 3% B at 0.1 min, and then increased linearly to 40% B at 100.9 min. Solvent B was increased to 95% at 101.0 min and maintained at 95% until 108.9 min. The gradient was returned to 0% B at 109.0 min and maintained at 0% B until 115.9 min, followed by post-run equilibration until 120 min.

#### HEK293T coimmunoprecipitation

Bound fraction peptides were analyzed on a Vanquish Neo nano-LC system coupled to an Orbitrap Exploris 240 mass spectrometer. A trap-and-elute configuration was used with an Acclaim PepMap 100 trapping column and an IonOpticks Aurora C18 analytical column (75 µm internal diameter × 15 cm). Mobile phase A was 5% acetonitrile/0.1% formic acid and mobile phase B was 80% acetonitrile/0.1% formic acid. Peptides were separated at 300 nL/min over a 60-min gradient: 0-10% B (0-5 min), 10–40% B (5-40 min), 40-100% B (40-42 min), held at 100% B (42-50 min), returned to 0% B (50–52 min), followed by post-run equilibration until 60 min.

For both methods, data were acquired in positive-ion mode using data-dependent acquisition. Full MS spectra were acquired in the Orbitrap at 120,000 resolution with monoisotopic precursor selection enabled in peptide mode. Precursors with charge states 2–5 and intensity ≥5,000 were selected for fragmentation; undetermined charge states were excluded, and isotope exclusion was enabled. MS/MS spectra were acquired at 15,000 resolution with a first mass of *m/z* 200 and centroid mode, using 30% normalized collision energy.

For proteomics, a 2-s cycle time was used with an *m/z* range of 350–1,600, automatic injection time, RF lens 50%, 30-s dynamic exclusion (±10 ppm), and a 1.4-*m/z* isolation window. For coimmunoprecipitation, up to 20 precursors were selected per full MS scan over *m/z* 400–1,600, with a 25-ms injection time, RF lens 60%, 30-s dynamic exclusion (±5 ppm), and a 1.5-*m/z* isolation window. MS/MS AGC targets were Standard setting with automatic injection time for proteomics and 100% (1 × 10⁵ ions) with a 50-ms injection time for coimmunoprecipitation.

#### Raw data processing

Raw global and phosphopeptide-enriched proteomics and coimmunoprecipitation data were processed using FragPipe (v23.0) against the human SwissProt database. Variable modifications included lysine and N-terminal acetylation (+42.0106 Da) and methionine oxidation (+15.9949 Da), along with serine, threonine and tyrosine phosphorylation (+79.96633) for phosphopeptide-enriched samples, allowing up to three variable modifications per peptide. DDA+ mode, MSBooster and Ion Quant were enabled, and peptide scoring was conducted via the Koina server (https://koina.wilhelmlab.org:443/v2/models/). Potential contaminants were removed from detected proteins. MaxLFQ intensities for global and phosphopeptide-enriched proteomics samples were log2-transformed and median-normalized.

### Bioinformatics

#### Principal component analysis

Log2-normalized MaxLFQ intensities for all biological replicates were compiled independently for the global and phosphopeptide-enriched proteomics datasets. Missing values were imputed using quantile regression imputation of left-censored data method using the “imputeLCMD” package (v2.1) in R (4.5.1) and RStudio (v4.5.1). For visualization purposes, batch correction was then performed using the “limma” package (v3.66.0), using differentiation group as the batch variable. Principal component analysis was run on the resulting matrices using the “PCAtools” package (v2.22.4) and visualized using the biplot function.

#### Data exploration

Log2-normalized MaxLFQ intensities were analyzed as three independent pairwise comparisons. Within each comparison, proteins/phosphopeptides were “flagged” if not present in at least 50% of samples. If a protein/phosphopeptide was “flagged” in both genotypes, it was filtered out of the matrix. Remaining data were separate by genotype and imputed using the “MSnbase” package (v2.36.0). Missing values for proteins/phosphopeptides detected in at least 50% of samples were considered missing at random and imputed using K-nearest neighbors; those detected in less than 50% of samples were considered missing not at random and imputed using Minimum Probability (49). Imputed values were merged with the original matrices to preserve already existing values.

#### Design model and differential abundance analysis

Each pairwise comparison was analyzed independently using “limma” with genotype as the primary variable and differentiation group as the variance group and blocking factor. Empirical Bayes moderation was applied, and p-values were adjusted using the Benjamini-Hochberg method. The “EnhancedVolcano” package (v1.28.2) was used to visualize differentially abundant proteins/phosphopeptides with p-value cutoff of 0.05 and log2-fold change (Log2FC) cutoff of 0.5.

#### Overlap analyses

Venn Diagrams generated for analyzing overlap in global and phosphopeptide-enriched proteomics data, along with coimmunoprecipitation candidate interactors and their disease associations, were generated using R “VennDiagram” (v1.8.2).

#### Gene Ontology

DAPs for all global proteomics pairwise comparisons were grouped by direction (increased/decreased abundance relative to comparison genotype). Metascape (v3.5.20260701) and/or SynGO (v1.3) were used to test for biological process enrichment in each group using Fisher’s exact test and false discovery rate correction. Statistically significant biological processes from both platforms were merged (-LogP > 2). GO term redundancy was simplified using Revigo (revigo.irb.hr) with similarity threshold of 0.5 (50). Terms were reduced to parent. Dot plots were visualized in using package “ggplot2” (v4.0.3). This process was repeated for coIP candidate interactors. Donut plots were visualized using “webr” (v0.1.5).

#### Gene-set enrichment analysis

Targeted GSEA was performed using the “fgsea” package (v1.36.2) to identify whether biological processes identified in the PACS1^(+/R203W)^ vs PACS1^(+/+)^ comparison were recapitulated in male and female patient-derived iNs. For each patient, proteins were ranked by Log2FC relative to their corresponding isogenic control. Proteins with NAs or duplicate symbols were excluded. Gene sets corresponding to biological processes in Figure 2D were queried using package “org.Hs.eg.db” (v3.22.0). GSEA was performed independently for each patient, with a gene set size 2-500. Normalized enrichment scores (NES) and Benjamini-Hochberg adjusted p-values were calculated for each GO term. NES values were visualized using “ggplot2”.

#### Kinase-substrate enrichment analysis

KSEA was performed for all pairwise comparisons using limma output from phosphopeptide-enriched proteomics samples using the “KSEAapp” package (v2.0) with default NetworKIN cutoff of 5 (38). Kinases with more than 2 substrates and Benjamini-Hochberg adjusted p-value ≤ 0.05 were considered true discoveries. Heatmaps were generated using “ggplot2”.

#### Kinase abundance in isogenic and patient lines

Kinase abundance was assessed across all biological replicates for each genotype. The average log2-normalized MaxLFQ intensity of each kinase in PACS1^(+/+)^ samples was used as the reference to calculate Log2FC for each sample. The same approach was applied to male and female patient-derived iNs using their corresponding isogenic controls to calculate the Log2FC. Heatmaps were generated using the “pheatmap” package (v1.0.13).

#### Phosphopeptide abundance in patient lines

Male and female PACS1 syndrome patient-derived iN phosphopeptide-enriched data were used to determine whether inferred differences in AKT1, CAMK2A, CSNK2A1, and MAP14 activity were recapitulated in patient samples. Substrates identified in our original analysis that are downstream of these kinases were evaluated in patient datasets. Log2FC were calculated by comparing log2-normalized MaxLFQ intensities in each patient to that of their corresponding isogenic control.

Candidate substrates were restricted to phosphopeptide annotated in PhosphoSitePlus with an absolute Log2FC > 1. Phosphopeptides were grouped by associated kinase. Log2FC in patient samples were visualized using the “pheatmap” package.

#### HEK293T coimmunoprecipitation analysis

Proteins in bound fractions of GFP, PACS1(R203)-GFP and PACS1(W203)-GFP samples were identified in 3 biological replicates. Proteins detected in at least 1 replicate were considered candidate interactions.

#### AlphaPulldown

The AlphaPulldown pipeline was used to predict protein-protein interactions between PACS1(R203)-GFP, PACS1(W203)-GFP, or GFP (bait) and 5,664 candidate (prey) proteins from iN global proteomics dataset (44). Multiple sequence alignments (MSAs) and template searches were performed on all bait and prey to generate features (proteins lacking features were excluded). AlphaPulldown “pulldown” mode was used to run AlphaFold-Multimer on each bait-prey pair. Predictions were analyzed using AlphaPulldown’s results table. Prey with an interface predicted template modeling score (iPTM) >0.6 and average interface predicted aligned error (PAE) <20 were deemed potential candidate interactors.

#### Candidate interactor analysis

Candidate interactors from *in silico* coIP and HEK293T coIP were cross-referenced against the Online Mendelian Inheritance in Man (OMIM) and Simons Foundation Autism Research Initiative (SFARI) Gene databases (October 2025) to identify proteins associated with NDDs, ASD, or other diseases.

### Electrophysiology

iNs were co-cultured with rat primary astrocytes and patched between d35-42 post doxycycline treatment. Assays were conducted using an upright Olympus BX51 microscope with epifluorescence capabilities and a SutterPatch Integrated Patch Amplifier (Sutter Instruments). A Sutter Puller P-2000 created the recording electrodes from borosilicate glass (resistance 2-4MΩ). The internal solution used in these electrodes was 120mM K-Gluc, 10mM Na-Gluc, 10mM K-HEPES, 4mM NaCl, 10 mM Na-Creatine, 4mM MgATP, 2mM NaATP, and 0.3mM NaGTP which was adjusted to 295mOsm and pH 7.4. Patched neurons were filled with Alexa-Fluor488 (Thermo Scientific, 1mM). The cells were immersed in ACSF bath solution (125 mM NaCl, 2.5 mM KCl, 2 mM CaCl_2_, 1 mM MgCl_2_, 1.25 mM NaH_2_PO4, 26 mM NaHCO_3_, 15 mM glucose, 1 mM myo-inositol, 2 mM Na-pyruvate, 0.4 mM ascorbic acid) which was oxygenated during experiments with 95%O_2_ and 5%CO_2_. Bath solution was heated to 33°C in the recording chamber.

Voltage clamp recordings of 500ms membrane step depolarizations starting at negative potentials (-90 or -75mV) and ending at positive potentials (+60 or +30mV) in 5mV steps. Sodium current was measured as the peak (minimum) of the response and steady-state potassium current was calculated as the mean of the last 10ms of the response. Action potential kinetics and spiking were analyzed in current clamp mode. Abrupt 40ms current injections increasing in magnitude (2 pA steps) were used until the first AP firing was achieved. Action potential measurement used the previously described parameters (51). For spiking analysis, 500ms current injections starting at - 10pA and ending at +180pA in 10pA steps were used and the evoked firing frequency was calculated.

### Statistics

Statistical analyses were performed using GraphPad Prism (v11.0.0) and R; tests specified in figure legends. Comparisons among PACS1^(+/+)^, PACS1^(+/R203W)^ and PACS1^(-/-)^ (or male patient vs female patient vs control) used Kruskal-Wallis tests with Dunn’s multiple comparisons. Kruskal-Wallis p-values are annotated p_Kruskal_; Dunn’s p-values for PACS1^(+/R203W)^ and PACS1^(-/-)^ versus PACS1^(+/+)^ are denoted p_R203W_ and p_KO_, respectively; Dunn’s p-values for male and female patient versus control are denoted as p_male_ and p_female_, respectively. Comparisons among two genotypes used 2-tailed Mann-Whitney tests. Two-way ANOVAs with Tukey’s multiple comparisons were used for Sholl analysis and action potential frequency vs. current injection. Bioinformatics p-values were adjusted using Benjamini-Hochberg method. Statistical significance was defined as p ≤ 0.05. Cell number was used as n for immunofluorescence and electrophysiology analyses. Biological replicates were used as n for all other analyses. Box plots show median, interquartile range, and minimum-to-maximum range. Bar plots show mean ± SEM.

### Study approval

This research was approved by the Northwestern University Institutional Review Board (IRB STU00215054) and followed ethical regulations relevant to the use of induced pluripotent stem cell lines.

## Data availability

All data and code are available upon request. Data used to generate figures, tables, and supplemental figures are provided in the Supporting Data Values file.

## AUTHOR CONTRIBUTIONS

This work was conceptualized by ALS and AGG. ALS performed all cell culture, wet lab experiments, microscopy and image analysis. The Northwestern Proteomics Core Facility prepared and analyzed global and phosphopeptide-enriched proteomics and HEK293T coimmunoprecipitation samples. ALS performed all bioinformatics analysis with guidance from XGM. SRG performed and analyzed electrophysiology experiments. CMP modified and ran AlphaPulldown pipeline. ALS generated all figures and wrote the manuscript, with edits by AGG. All authors read and approved the final manuscript.

## FUNDING SUPPORT

This work was funded by the PACS1 Syndrome Research Foundation (AGG), National Institutes of Health (NINDS R01NS123163 [AGG], R01NS140046 and DP2NS148744 [RSS], NIGMS T32-GM008061 [ALS] and 1F31GM151811-01[CMP]), the National Science Foundation Synthetic Biology Across Scale National Research Traineeship (DGE2021900 [CMP]), and Northwestern University Driskill Graduate Program in Life Sciences (ALS, CMP).

## ACKNOWLEDGMENTS

Lentiviruses were generated by the Gene Editing, Transduction, and Nanotechnology (GET iN) Core (RRID:SCR_026417) through the Northwestern University Skin Biology & Diseases Resource-Based Center (NIH P30AR075049). Proteomics services were performed by the Northwestern Proteomics Core Facility (RRID:SCR_017945), funded by NCI CCSG P30 CA060553 to the Robert H Lurie Comprehensive Cancer Center and NIGMS RM1 GM156535 to the National Center for Translational and Developmental Proteomics. Microscopy was performed on a Nikon CSU-W1 SoRa system (1S10OD032270-01) at the Northwestern University Center for Advanced Microscopy (RRID:SCR_020996), supported by NCI CCSG P30 CA060553. Computational resources and staff support were provided by Northwestern University’s Quest high-performance computing facility, supported by the Office of the Provost, the Office for Research, and Northwestern University Information Technology.

We thank Gemma Carvill, Scott Adney, and their respective labs for their continued support and input. We thank Peter Faull, Szymon Filip, and Aarohan Mattamana from the Northwestern Proteomics Core Facility for their guidance and technical support. ALS also thanks Yvette Wong, Erica Davis, and Brian Mitchell for their feedback.

## Conflict of Interest Statement

The authors have declared that no conflict of interest exists.

## Notes

### Competing Interest Statement

The authors have declared no competing interest.

