## Supplement for "Molecular and functional profiling distinguishes PACS1 syndrome variant from PACS1 loss-of-function in iNeurons"

**This PDF file includes:**

**Figure S1-S2, Tables S1-S6**

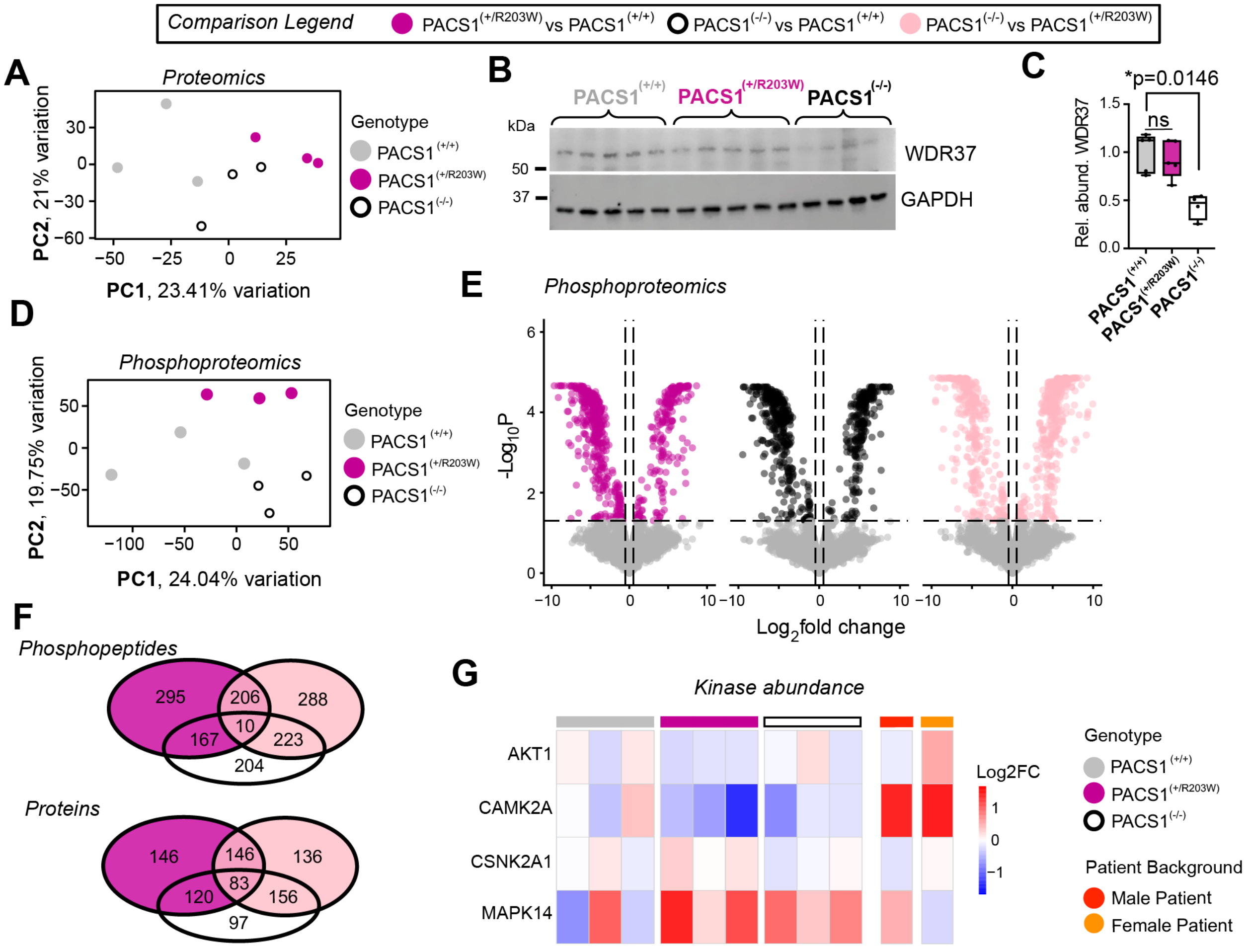


**Figure S1. Supporting analyses for global and phosphopeptide-enriched proteomic profiling** (corresponding to Figure 2). **(A)** Principal component analysis of imputed and batch-corrected global proteomic samples from d29 iNs. **(B)** Validation Western blot for WDR37 and GAPDH protein abundance in d29 iNs. **(C)** Quantification of the ratio of WDR37:GAPDH in d29 PACS1$^{(+/+)}$, PACS1$^{(+/R203W)}$ and PACS1$^{(-/-)}$ iNs (Kruskal-Wallis with Dunn's multiple comparisons; $p_{Kruskal}$=0.0050; $p_{R203W}$>0.9999, $p_{KO}$=0.0146; n=5 samples for PACS1$^{(+/+)}$, PACS1$^{(+/R203W)}$ and n=4 samples for PACS1$^{(-/-)}$ iNs). **(D)** Principal component analysis of imputed and batch-corrected phosphopeptide-enriched proteomic profiles from d29 iNs. **(E)** Volcano plot of statistically significant DAPhs with an absolute Log2FC > 0.5 across pairwise genotype comparisons (p-values≤ 0.05; n= 8502, 8601, 8476 phosphopeptides for PACS1$^{(+/R203W)}$ vs PACS1$^{(+/+)}$, PACS1$^{(-/-)}$ vs PACS1$^{(+/+)}$, PACS1$^{(-/-)}$ vs PACS1$^{(+/R203W)}$ comparisons, respectively). **(F)** Venn diagrams showing the overlap of DAPhs and their associated proteins across pairwise genotype comparisons. **(G)** Abundance (represented as Log2FC) of kinases AKT1, CAMK2A, CSKK2A1, and MAPK14 in the global proteomics dataset for all PACS1$^{(+/+)}$, PACS1$^{(+/R203W)}$ and PACS1$^{(-/-)}$ (Log2FC calculated using the average intensity of each kinase in PACS1$^{(+/+)}$ samples), and male patient and female patient (Log2FC calculated using the intensity of each kinase in their respective isogenic control sample).

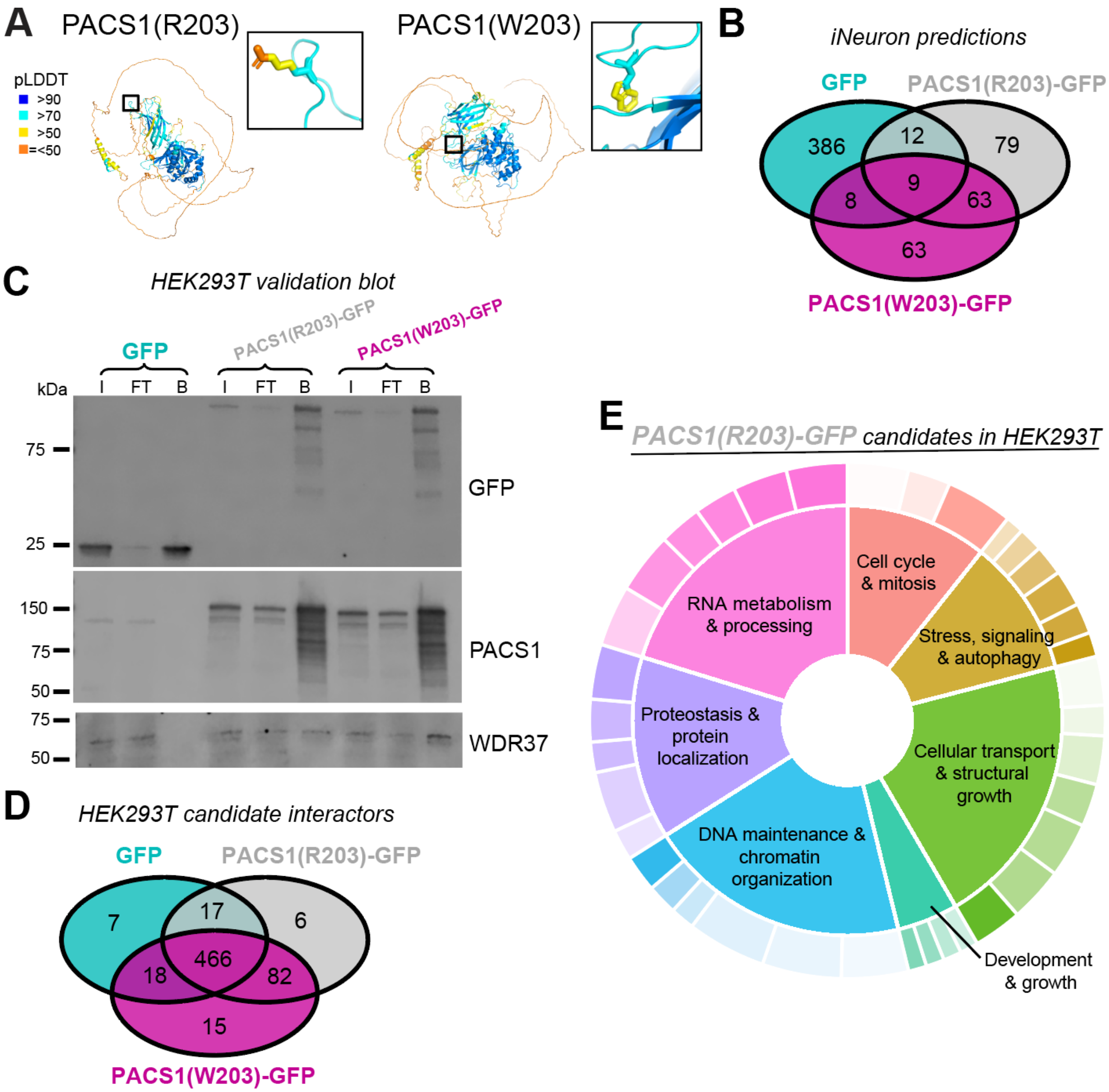


**<u>Figure S2. Supporting analyses for PACS1 comparative coimmunoprecipitation</u>** (corresponding to Figure 5). **(A)** Confidence scores for AlphaFold3 structural models of PACS1 containing either R203 or W203 residue. **(B)** Overlap of predicted candidate interactors for GFP, PACS1(R203)-GFP and PACS1(W203)-GFP, identified by *in silico* coIP. **(C)** Western blot validation of coIP in HEK293T demonstrating blot against GFP, PACS1 and positive control WDR37 in Input (I), Flow-through (FT) and Bound (B) fractions. **(D)** Overlap of candidate interactors for GFP, PACS1(R203)-GFP and PACS1(W203)-GFP, identified by coIP in HEK293T cells. **(E)** Gene ontology analysis of HEK293T candidate interactors for PACS1(R203)-GFP, grouped by shared molecular function; sizing of pie slice dependent on gene count for that GO-term/group. All HEK293T candidate interactors listed in Table S6.

**Table S1. List of top 15 DAPs identified in global proteomics, unique to each pairwise comparison**

| Comparison | Protein | UniprotID | Log2FC | p-value |
|---|---|---|---|---|
| PACS1$^{(+/R203W)}$ vs PACS1$^{(+/+)}$ | FJX1 | Q86VR8 | -6.6286168 | 2.47E-05 |
| | TRIM9 | Q9C026 | -6.5186037 | 2.47E-05 |
| | SSC5D | A1L4H1 | -6.1914176 | 2.47E-05 |
| | JAK1 | P23458 | -5.4672678 | 2.47E-05 |
| | GBP1 | P32455 | -5.4198556 | 2.47E-05 |
| | SLC8A3 | P57103 | -5.4190983 | 2.47E-05 |
| | TNS1 | Q9HBL0 | -5.3843418 | 2.47E-05 |
| | ITGA1 | P56199 | -5.2127681 | 2.47E-05 |
| | GCAT | O75600 | -5.174398 | 2.47E-05 |
| | WIPF3 | A6NGB9 | -5.0098119 | 2.47E-05 |
| | CAPG | P40121 | -4.6959291 | 2.47E-05 |
| | KCTD10 | Q9H3F6 | 5.05061989 | 2.47E-05 |
| | ASH2L | Q9UBL3 | 5.29644226 | 2.47E-05 |
| | MCM2 | P49736 | 6.21660005 | 2.47E-05 |
| | PPP2R3A | Q06190 | 6.55347469 | 2.47E-05 |
| PACS1$^{(-/-)}$ vs PACS1$^{(+/+)}$ | PACS1 | Q6VY07 | -8.0460917 | 4.22E-09 |
| | ADGB | Q8N7X0 | 6.37139479 | 1.14E-08 |
| | MTMR4 | Q9NYA4 | 9.43332236 | 1.04E-08 |
| | EFNB1 | P98172 | 7.65310314 | 1.60E-08 |
| | FJX1 | Q86VR8 | -6.9829145 | 6.71E-08 |
| | BECN1 | Q14457 | -6.2948409 | 4.68E-08 |
| | LMBRD1 | Q9NUN5 | -6.2026164 | 6.53E-08 |
| | LYPD1 | Q8N2G4 | -6.1577241 | 6.43E-08 |
| | TMEM222 | Q9H0R3 | -6.0365097 | 2.76E-08 |
| | SLC6A17 | Q9H1V8 | -5.6949542 | 5.42E-08 |
| | CD55 | P08174 | -5.6148888 | 7.52E-08 |
| | NACC1 | Q96RE7 | -4.6784015 | 4.20E-08 |
| | CMTR1 | Q8N1G2 | 5.56418552 | 7.70E-08 |
| | NUMB | P49757 | 6.22872355 | 5.15E-08 |
| | CRYAA | P02489 | 6.72550779 | 7.58E-08 |
| PACS1$^{(-/-)}$ vs PACS1$^{(+/R203W)}$ | PACS1 | Q6VY07 | -8.1174107 | 4.57E-09 |
| | TRIM9 | Q9C026 | 7.00008593 | 6.64E-09 |
| | PLAC9 | Q5JTB6 | 7.09188607 | 3.61E-09 |
| | MED12L | Q86YW9 | 7.54928274 | 6.95E-09 |
| | EBAG9 | O00559 | 8.97536774 | 5.21E-09 |
| | STX17 | P56962 | 5.33658509 | 1.23E-08 |
| | VARS2 | Q5ST30 | 6.65988246 | 9.98E-09 |
| | MAZ | P56270 | 7.24087906 | 1.10E-08 |
| | RABIF | P47224 | -6.2474855 | 1.40E-08 |
| | RGPD3 | A6NKT7 | -6.1208962 | 2.41E-08 |
| | PELI2 | Q9HAT8 | -4.9025403 | 2.14E-08 |
| | SRBD1 | Q8N5C6 | 6.10942894 | 2.42E-08 |
| | NMNAT2 | Q9BZQ4 | 7.04380769 | 2.31E-08 |
| | GCAT | O75600 | 6.37403597 | 2.86E-08 |
| | LRRN2 | O75325 | 5.29150581 | 3.55E-08 |

DAPs were ranked by statistical significance (adjusted p-value) and Log2FC.

**Table S2. List of top 10 GO terms identified in global proteomics, unique to each pairwise comparison**

| Comparison | GO TermID | GO Name | -LOG(P) | Gene Count | Direction |
|---|---|---|---|---|---|
| PACS1$^{(+/R203W)}$ vs PACS1$^{(+/+)}$ | GO:0006364 | rRNA processing | 6.48 | 8 | UP |
| | GO:0045766 | positive regulation of angiogenesis | 4.91 | 7 | DOWN |
| | GO:0051052 | regulation of DNA metabolic process | 3.74 | 8 | UP |
| | GO:0010810 | regulation of cell-substrate adhesion | 3.70 | 6 | DOWN |
| | GO:0045596 | negative regulation of cell differentiation | 3.68 | 9 | UP |
| | GO:0006259 | DNA metabolic process | 3.46 | 9 | UP |
| | GO:0098609 | cell-cell adhesion | 3.44 | 9 | DOWN |
| | GO:0006399 | tRNA metabolic process | 3.26 | 5 | UP |
| | GO:0016567 | protein ubiquitination | 3.14 | 8 | UP |
| | GO:0001944 | vasculature development | 3.13 | 6 | UP |
| PACS1$^{(-/-)}$ vs PACS1$^{(+/+)}$ | GO:0009451 | RNA modification | 7.42 | 9 | UP |
| | GO:0006400 | tRNA modification | 6.92 | 7 | UP |
| | GO:0060261 | positive regulation of transcription initiation by RNA polymerase II | 5.36 | 5 | UP |
| | GO:0043414 | macromolecule methylation | 5.18 | 6 | UP |
| | GO:0032259 | methylation | 4.95 | 6 | UP |
| | GO:2000142 | regulation of DNA-templated transcription initiation | 4.58 | 5 | UP |
| | GO:0018193 | peptidyl-amino acid modification | 4.31 | 6 | DOWN |
| | GO:0007034 | vacuolar transport | 4.18 | 5 | DOWN |
| | GO:0050803 | regulation of synapse structure or activity | 3.94 | 6 | DOWN |
| | GO:1902903 | regulation of supramolecular fiber organization | 3.64 | 6 | DOWN |
| PACS1$^{(-/-)}$ vs PACS1$^{(+/R203W)}$ | GO:0006367 | transcription initiation at RNA polymerase II promoter | 4.60 | 7 | UP |
| | GO:0050767 | regulation of neurogenesis | 4.05 | 6 | DOWN |
| | GO:0032535 | regulation of cellular component size | 4.05 | 6 | DOWN |
| | GO:0090066 | regulation of anatomical structure size | 3.67 | 9 | UP |
| | GO:0043122 | regulation of canonical NF-kappaB signal transduction | 3.64 | 8 | UP |
| | GO:0007281 | germ cell development | 3.30 | 7 | UP |
| | GO:0061025 | membrane fusion | 3.26 | 5 | UP |
| | GO:0032535 | regulation of cellular component size | 3.26 | 7 | UP |
| | GO:0007167 | enzyme-linked receptor protein signaling pathway | 3.25 | 7 | DOWN |
| | GO:0090066 | regulation of anatomical structure size | 3.20 | 6 | DOWN |

GO terms were ranked by statistical significance (-LogP) and gene count and represent parent terms with reduced redundancy. GO terms were identified as enriched or diminished based on proteins identified as DAPs in each global proteomics pairwise comparison.

**Table S3. Electrophysiology analysis and statistics of isogenic PACS1$^{(+/+)}$, PACS1$^{(+/R203W)}$ and PACS1$^{(-/-)}$ iNs**

| | PACS1$^{(+/+)}$ | PACS1$^{(+/R203W)}$ | PACS1$^{(-/-)}$ | P value (Kruskal-Wallis) | Posthoc (vs. PACS1$^{(+/+)}$) | Figure |
|---|---|---|---|---|---|---|
| n | 12-16 | 13-16 | 16-18 | *NA* | *NA* | *NA* |
| **AHP (mV)** | -40.41 ±1.872 | -39.91 ±1.928 | -36.96 ±1.97 | 0.4629 | (+/R203W) >0.9999<br>(-/-) = 0.5651 | *NA* |
| **Threshold (mV)** | -36.6 ±1.250 | -36.13 ±0.692 | -38.45 ±1.471 | 0.5622 | (+/R203W) >0.9999<br>(-/-) = 0.8468 | *NA* |
| **Rheobase (pA)** | 108.6 ±8.908 | 96.35 ±10.50 | 98.3 ±9.891 | 0.6264 | (+/R203W) = 0.6822<br>(-/-) > 0.9999 | 4E |
| **Amplitude (mV)** | 58.27 ±3.003 | 59.94 ±2.158 | 60.86 ±2.543 | 0.6658 | (+/R203W) >0.9999<br>(-/-) = 0.7480 | 4F |
| **Halfwidth (ms)** | 1.946 ±0.0939 | 1.972 ±0.152 | 2.021 ±0.1694 | 0.9871 | (+/R203W) >0.9999<br>(-/-) > 0.9999 | 4G |
| **Max K (pA/pF)** | 82.20 ±8.973 | 83.23 ±13.83 | 86.40 ±10.60 | 0.8976 | (+/R203W) >0.9999<br>(-/-) > 0.9999 | *NA* |
| **Max Na (pA/pF)** | -69.23 ±8.692 | -65.14 ±11.92 | -76.72 ±9.048 | 0.4518 | (+/R203W) >0.9999<br>(-/-) > 0.9999 | *NA* |
| **Resistance (μΩ)** | 0.5817 ±0.0662 | 0.6669 ±0.0933 | 0.5339 ±0.06952 | 0.5148 | (+/R203W) >0.9999<br>(-/-) = 0.9380 | *NA* |
| **Capacitance (pF)** | 53.68 ±3.898 | 56.68 ±4.938 | 51.27 ±4.252 | 0.7370 | (+/R203W) >0.9999<br>(-/-) > 0.9999 | *NA* |

N indicates the number of cells assayed. Max K and Max Na are capacitance-adjusted. Data is presented as the mean ± SEM. Data were analyzed using Kruskal-Wallis test followed by Dunn's multiple comparisons test for PACS1$^{(+/R203W)}$ and PACS1$^{(-/-)}$ iNs in comparison to PACS1$^{(+/+)}$ control iNs. All p-values displayed in table. Significance defined as $p \leq 0.05$.

**Table S4. Electrophysiology analysis and statistics of iNs derived from male and female PACS1 syndrome patients compared to summary control line**

| | Control | Male Patient | Female Patient | P value (Kruskal-Wallis) | Posthoc (vs. Control) | Figure |
|---|---|---|---|---|---|---|
| **n** | 8 | 8 | 11 | *NA* | *NA* | *NA* |
| **AHP (mV)** | -15.81 ±0.8440 | -18.58 ±1.538 | -18.07 ±1.116 | 0.3203 | male =0.4916<br>female = 0.3108 | *NA* |
| **Threshold (mV)** | -39.91 ±0.6503 | -40.17 ±1.165 | -39.94 ±0.9194 | 0.9530 | male >0.9999<br>female >0.9999 | *NA* |
| **Rheobase (pA)** | 34.18 ±7.323 | 41.50 ±8.080 | 32.02 ±4.919 | 0.5331 | male = 0.6396<br>female > 0.9999 | 4I |
| **Amplitude (mV)** | 55.70 ±2.996 | 82.63 ±3.045 | 77.88 ±3.716 | * 0.0008 | **male =0.0010<br>**female = 0.0047 | 4J |
| **Halfwidth (ms)** | 2.392 ±0.3498 | 0.8258 ±0.04359 | 1.125 ±0.08468 | ****<0.0001 | ****male <0.0001<br>**female = 0.0098 | 4K |
| **Max K (pA/pF)** | 36.31 ±6.544 | 56.17 ±8.175 | 41.70 ±5.802 | 0.1924 | male = 0.1663<br>female > 0.9999 | *NA* |
| **Max Na (pA/pF)** | -69.80 ±13.39 | -92.59 ±18.60 | -74.36.80 ±11.55 | 0.5565 | male = 0.6269<br>female > 0.9999 | *NA* |
| **Resistance (μΩ)** | 1.385 ±0.4090 | 1.027 ±0.1419 | 0.9024 ±0.1115 | 0.9402 | male >0.9999<br>female >0.9999 | *NA* |
| **Capacitance (pF)** | 41.53 ±5.031 | 60.67 ±5.117 | 47.46 ±5.076 | 0.0632 | * male = 0.0395<br>female = 0.6302 | *NA* |

N indicates the number of cells assayed. Max K and Max Na are capacitance-adjusted. Data is presented as the mean ± SEM. Data were analyzed using Kruskal-Wallis test followed by Dunn's multiple comparisons test for male and female patient iNs in comparison to summary line control. All p-values displayed in table. Significance defined as $p \leq 0.05$.

**Table S5. Lists of predicted candidate interactors in iNs unique to or shared by PACS1(R203)-GFP and PACS1(W203)-GFP**

| **PACS1(R203)-GFP** (79) | | | | **PACS1(W203)-GFP** (63) | | | **Shared** (63) | | |
|---|---|---|---|---|---|---|---|---|---|
| HIF1AN | MRPL53 | RAB2B | CHID1 | SRP72 | TUBA1C | EXOSC7 | HSPA1A | RHOC | RAB13 |
| RAP2A | DCAF7 | GAPDH | P4HA1 | GOLPH3 | SSU72 | HPRT1 | RPL8 | UBE2L6 | MRPS22 |
| NRAS | AP1S2 | UNC119 | SCAI | OSGEPL1 | RPL23 | PDXP | HRAS | EMC8 | SAE1 |
| MRPL22 | SLC35E1 | SUCLG1 | PITHD1 | GLIPR2 | MAGOHB | TSFM | EXTL3 | LAMTOR4 | HDAC1 |
| ACOT8 | KDM1A | MMUT | RAB8A | RAB3D | EXOSC2 | RHOA | URM1 | NUP37 | BUB3 |
| SNRNP40 | PPP2CA | PMVK | ITFG1 | PTS | RAB10 | MRPL44 | TUBB1 | RPS8 | EXOG |
| IFT25 | CAB39 | RPS29 | CLVS1 | PNO1 | THOC6 | FKBP7 | NCBP2 | ARMH3 | TRUB1 |
| FBXL2 | SEC11A | TMEM147 | POP7 | TUBB4A | TUBAL3 | RPS21 | ATP6V0D1 | CAPZB | GLTP |
| CTSA | RPF2 | HDAC2 | MRPL14 | GPD1L | MLST8 | COPZ1 | ATP5F1A | ATG5 | COPS8 |
| SDHAF2 | AKR1C2 | PAAF1 | PLD3 | LRRC8A | NIP7 | DERL2 | CBFB | PPIL1 | RAN |
| POLR2J | SMYD3 | DPM1 | TIMP3 | RPN1 | RCC1 | CTU1 | WDR45B | CIAO2B | SNX12 |
| GSPT1 | ATRAID | PFKL | C17orf75 | MRPS25 | OSTM1 | GGT7 | SUMO1 | RPS14 | RAB2A |
| EMC2 | RHOT1 | WDR48 | MRPL19 | GANAB | TRMT61A | EXOSC5 | NDUFS8 | PCNA | RSU1 |
| HEXB | CYBC1 | MOCS2 | POLR2L | CSNK1A1 | FAM91A1 | PAFAH1B3 | ORMDL1 | EIF4E3 | SOD1 |
| PSMD2 | RALA | CNPY2 | PGAM5 | PPP2CB | RPE | SH3BGRL2 | EIF5A | DNAAF10 | METTL1 |
| PFN1 | GLB1 | NPLOC4 | DIABLO | RAB32 | CYB5R1 | MRPL23 | NDUFV1 | AGPAT5 | RPS28 |
| MAT2A | PTCD3 | WDR12 | | PITPNB | GAN | GMPS | LIPA | GGACT | UBE2N |
| GTF2A2 | RAB22A | ALDH16A1 | | TPBG | DPP10 | EIF2B1 | SUPT4H1 | TRAPPC3 | COX15 |
| TBCK | NCS1 | ARL5A | | RANGRF | ATP5F1B | DIRAS2 | EIF4A3 | CNOT10 | NAA50 |
| TFRC | NUP43 | ARF4 | | AKR1C3 | TBC1D23 | PNCK | WDR77 | RAB7A | MTMR9 |
| CIAO1 | VPS26C | MAN2B2 | | WDR6 | STX1B | DMAC2L | SH3BGRL | DIRAS1 | ARL2 |

Candidate interactors were identified through an *in silico* coimmunoprecipitation using the iNeuron proteome and the AlphaPulldown pipeline. Proteins used for this analysis were filtered based on iPTM > 0.6 and average interface PAE < 20. Proteins predicted to interact with the GFP only control were excluded from this analysis.

**Table S6. Lists of predicted candidate interactors in HEK293T unique to or shared by PACS1(R203)-GFP and PACS1(W203)-GFP**

| **PACS1(R203)-GFP** (6) | **PACS1(W203)-GFP** (15) | **Shared** (82) | | | | | |
|---|---|---|---|---|---|---|---|
| GTPBP4 | VPS4A | DDX39A | TARBP1 | LSG1 | CPSF7 | CTC1 | HADHB |
| USP9X | DMAP1 | RHOC | SMARCB1 | WDR11 | ZNF598 | RNPS1 | HSPA2 |
| MRPL21 | RBM33 | YWHAQ | RPL18 | HELZ2 | PACS1 | KPNB1 | SUB1 |
| DHX57 | PTRH1 | PRRC2C | PPP1CA | SRRT | LARP1 | KIF22 | RAB5C |
| ATM | TRMT1L | WDR37 | USP46 | TUBB2B | SZT2 | RBM39 | FXR2 |
| DSTN | SF3B2 | RUVBL1 | DCAF7 | GRWD1 | CEP170 | GIT2 | RPL34 |
| | ASPH | MCM8 | SEC13 | RBM15 | CEP78 | ARHGEF7 | ATP5PO |
| | AFDN | SEH1L | SIPA1L2 | SGF29 | MAP7D1 | DHX34 | RANGAP1 |
| | RPL5 | MRPL24 | TAB2 | PLCG1 | SEC22B | NOL6 | PRKCI |
| | CNP | DPF2 | NHP2 | CCNB1 | OBSL1 | PRMT5 | HADHA |
| | PAPLN | PRPF31 | GNL3L | ACTL6A | DPM1 | KIF2A | HSPA9 |
| | NCOR1 | PARD3 | AVEN | BAG2 | EIF4E2 | VAPB | MYH10 |
| | RNF40 | PIK3C3 | NT5DC2 | ACSL3 | LANCL1 | AP2A2 | HSPA1L |
| | OPA1 | RBM15B | TDRD3 | PTBP1 | POLD1 | | |
| | NDUFS8 | | | | | | |

Candidate interactors were identified in the bound fractions of PACS1(R203)-GFP and PACS1(W203)-GFP coimmunoprecipitations, performed in HEK293T cells. Proteins identified in the bound fraction of the GFP only control were excluded from this analysis. A protein was classified as a candidate interactor if it was present in at least one of three biological replicates.
